# On the impact of contaminants on the accuracy of genome skimming and the effectiveness of exclusion read filters

**DOI:** 10.1101/831941

**Authors:** Eleonora Rachtman, Metin Balaban, Vineet Bafna, Siavash Mirarab

**Affiliations:** Bioinformatics and Systems Biology Graduate Program, UC San Diego, CA 92093, USA; Department of Computer Science and Engineering, UC San Diego, CA 92093, USA; Department of Electrical and Computer Engineering, UC San Diego, CA 92093, USA

**Keywords:** Genome skimming, Kraken, contamination, shotgun sequencing, filtering

## Abstract

The ability to detect the identity of a sample obtained from its environment is a cornerstone of molecular ecological research. Thanks to the falling price of shotgun sequencing, genome skimming, the acquisition of short reads spread across the genome at low coverage, is emerging as an alternative to traditional barcoding. By obtaining far more data across the whole genome, skimming has the promise to increase the precision of sample identification beyond traditional barcoding while keeping the costs manageable. While methods for assembly-free sample identification based on genome skims are now available, little is known about how these methods react to the presence of DNA from organisms other than the target species. In this paper, we show that the accuracy of distances computed between a pair of genome skims based on k-mer similarity can degrade dramatically if the skims include contaminant reads; i.e., any reads originating from other organisms. We establish a theoretical model of the impact of contamination. We then suggest and evaluate a solution to the contamination problem: Query reads in a genome skim against an extensive database of possible contaminants (e.g., all microbial organisms) and filter out any read that matches. We evaluate the effectiveness of this strategy when implemented using Kraken-II, in detailed analyses. Our results show substantial improvements in accuracy as a result of filtering but also point to limitations, including a need for relatively close matches in the contaminant database.

## 1 Introduction

Anthropogenic pressure and other natural causes have resulted in severe disruption of the global ecosystems in recent years, including loss of biodiversity and invasion of non-native flora and fauna. Conservationists, struggling with an unprecedented rate of extinction, are using innovative approaches to measure the changing biodiversity of the planet. Genome sequencing provides an attractive alternative to physical sampling and cataloging, as falling costs have made it possible to shotgun sequence a reference specimen sample for at most $10 per Gb (with another $60 for sample prep). However, the analysis typically requires assembling and finishing a reference genome, which can still be prohibitively costly. It could be many decades before the biodiversity of our planet is represented in the form of finished genomes (and cataloged genomic variants) and before biodiversity measurements for each population can be acquired on an ongoing basis.

The standard molecular technique for measuring biodiversity at the organismal level is barcoding (Hebert et al., 2003; Savolainen et al., 2005; Taberlet et al., 2012), which involves DNA sequencing of taxonomically informative and group-specific marker genes (e.g., mtDNA COI (Hebert et al., 2003; Seifert et al., 2007), 12S/16S (Vences et al., 2005), plastid genes (Hollingsworth et al., 2009), and ITS (Schoch et al., 2012)). Existing reference databases and computational methods enable measurements of biodiversity using barcodes (Ratnasingham and Hebert, 2007; Steinke et al., 2005; Taberlet et al., 2012). However, since barcodes are short regions, their phylogenetic signal is limited (Hickerson et al., 2006). For example, 896 of the 4,174 species of wasps could not be distinguished from other species using COI barcodes (Quicke et al., 2012).

As an alternative, a *genome skim* is a low-coverage acquisition of short reads from a sample, typically around 1-5 Gbp (Coissac et al., 2016; Dodsworth, 2015), providing 0.1-10×coverage, and usually insufficient for assembling nuclear contigs. Falling sequencing costs have made genome-skimming cost-effective while providing richer data than barcoding, but the data is harder to analyze. Skimming applications often rely on assembling organelle genomes (e.g., Malé et al., 2014; Weitemier et al., 2014) from their over-represented reads. This approach throws away the vast majority of the reads, potentially limiting the resolution. Moreover, organelle genomes may not represent the rest of the genome and are not always easy to assemble. Ideally, we should use both reads from both nuclear and organelle genomes. However, methods that seek to mine all information from genome skims must be *assembly-free* and *map-free* and face additional challenges.

Recently, Sarmashghi et al., 2019 developed a method, Skmer, that accurately computes genomic distance between genome skims by simply analyzing *k*-mers (short substrings of length *k*) in both genome skims. Skmer is based on three principles. First, as observed by Ondov et al. (2016), the *Jaccard index, J* (the size of the intersection of two sets divided by the size of their union) between *k-mer* sets of the two genomes can be computed efficiently. Second, *J* can be used to estimate the genomic distance (*D*) between two species by carefully accounting for dependence on coverage, sequencing error, and genome length. Third, both coverage and error rate can be computed from genome skim data by modelling histograms of *k*-mer frequencies. By combining these three principles, Skmer provides excellent accuracy in estimating distances between genome skims. These distances can then be used for taxonomic identification and phylogenetic placement (Balaban et al., 2019) of query genome skims with respect to a set of reference genome skims. Previous results have shown high accuracy and increased resolution compared to barcodes when using genome skims for taxonomic identification (Balaban et al., 2019; Sarmashghi et al., 2019).

The Skmer methodology, however, completely ignores the very real possibility that *a genome skim includes extraneous reads originating from other species*, often bacteria, virus or fungi, that cohabit inside the biological organism. With a slight abuse of terminology, we refer to all reads originating from species other than the target species being identified as *contamination*. Contamination of genome skims is unavoidable in many cases as microorganisms that co-exist with a species are often hard or impossible to separate from the original sample. To make matters worse, lab protocols used for genome skimming also can add human and other forms of contamination. The standard organelle-based analyses of skims manages to deal with sequencing errors and contamination by focusing on and assembling a small portion of the reads. These contaminates have the potential to mislead the Jaccard-based calculation of distance using methods such as Skmer. Thus, to take advantage of *all* reads across the genome, contaminants will have to be dealt with.

In this paper, we study the impact of contamination on Skmer estimates of the genomic distance. We then study whether the negative impact of contamination can be reduced using “exclusion filters”: search every read of a skim against a library of all known contaminants (e.g., bacterial, fungal, and viral genomes), filter out reads that map to the library, and use the remaining reads to compute the distance. The efficacy of this exclusion filtering approach is unclear and can depend on several factors, which we thoroughly explore here. We study these effects both based on a theoretical model and in careful simulation and empirical analyses using a leading read matching tool called Kraken-II (Wood et al., 2019).

## 2 Material and methods

### 2.1 Theoretical Exposition

Consider two genomes of equal length and separated by genomic distance *D*, defined as the portion of positions that do not match in a perfect alignment of the two genomes. Let *ρ* denote the proportion of *k*-mers in one species that are absent in the other. The Jaccard-index of *k*-mers is given by (Fig. 1a):

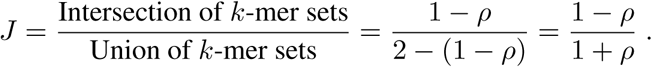

**Figure 1:**
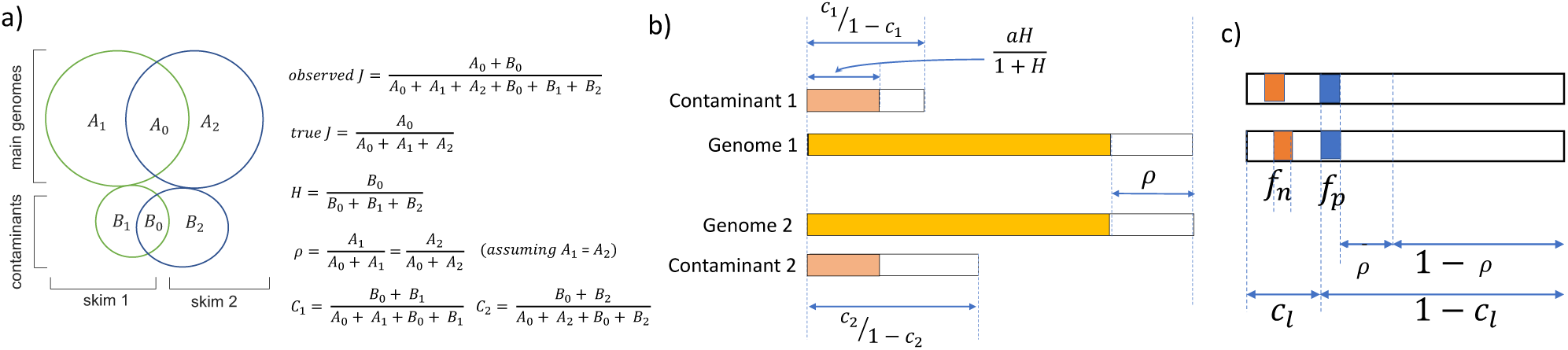
Model. (a) Definition of terms. Contaminating *k*-mers change the estimated Jaccard in a complex manner. (b) Assuming equal lengths for the two genomes, all quantities are measured as a fraction of the number of *k*-mers in each genome: 1 − *ρ* of the *k*-mers are shared between the base genomes; additionally, out of a total of 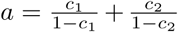 contaminants, 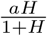 are shared, making the total number of shared *k*-mers equal to 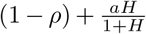 and the total number of distinct *k*-mers in the union equal 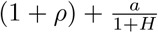. See Appendix A.1 for details. (c) Impact of false positives and negatives in contaminant removal in the disjoint contaminant scenario. We keep (1 − *c*_*l*_)(1 − *f*_*p*_) + *c*_*l*_*f*_*n*_ of the *k*-mers in each set, with the intersection proportion being (1 − *c*_*l*_)(1 − *f*_*p*_)(1 − *ρ*).

Assuming a uniform distribution of mutations along the genome, 𝔼(*ρ*) = 1 − (1 − *D*)^*k*^, and thus, we can estimate (Fan et al., 2015):

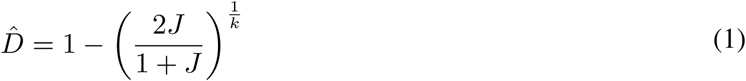

Skmer further models coverage and sequencing error and uses

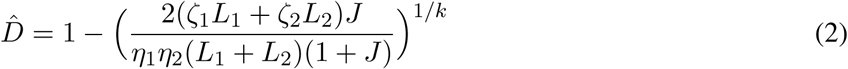

where *η*_*i*_, *ζ*_*i*_, and *L*_*i*_ are parameters related to coverage, error, and genome length, all automatically estimated by Skmer from *k*-mer profiles. As the simpler equation is easier to manipulate, we use (1) in our theoretical exposition. However, our empirical results will use the Skmer software, which uses (2). Throughout the paper, we use the relative error to quantify any error in estimating D:

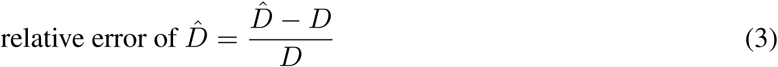

where *D* is the true genomic distance and 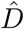 is the estimated genomic distance.

#### 2.1.1 Impact of contamination

Contamination can clearly alter Jaccard and hence the estimated genomic distance (Fig. 1a). The impact of contamination depends on factors such as the amount and exact composition of contaminants. For exposition purposes, let us assume that an identical proportion of *k*-mers (denoted by *c*_*l*_) of both skims are contaminated, and contaminant *k*-mers are entirely disjoint between the two genome skims. Then, *J* becomes a function of *c*_*l*_:

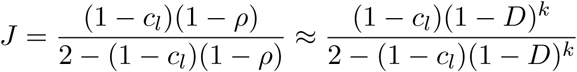

where the approximation is achieved by replacing *ρ* with its expectation.

Under these assumptions, Jaccard reduces under contamination and extent of reduction depends on *c*_*l*_ and to a lesser degree on *D* (Fig. S1a). If the impact of contamination on Jaccard is ignored, distance will be overestimated at a level that strongly depends on the true distance (Fig. S1b). When *D* is sufficiently high, substantial levels of contamination result in relatively low errors. However, with smaller distances, contamination can drastically increase the relative error. At *D* = 0.001 (e.g., within species differentiation), 3% contamination is enough to cause 100% relative error. Thus, under the simple disjoint contamination model, contamination has a large negative impacts *only* when the distance between base genomes is small.

Disjoint contamination assumptions, however, is quite strong. When both samples are contaminated with the same species (say, human), the assumption of disjoint contaminant *k*-mers can mislead. To generalize, consider two genomes with an equal number of *k*-mers *L*. Let *c*_1_ denote the fraction of the *k*-mers from sample 1 that are contaminated. Then, the ratio of contaminated *k*-mers to true *k*-mers in genome 1 is given by 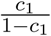 (Appendix A.1). Define *c*_2_ in an analogous fashion for genome 2, and let 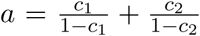 (Fig. 1a). Removing the disjoint contamination assumption, define the Jaccard index between the *k*-mers of the contaminants of the two samples as *H*. Then, as shown in Appendix A.1 and Figure 1b,

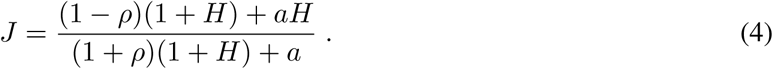

Plotting this formula shows that depending on *H*, the estimated Jaccard may over-estimate or under-estimate the true Jaccard, and converting the Jaccard to distance without any consideration of contaminants can lead to over or under-estimate the true distance (Fig. 2a). Once again, error depends on the true distance *D*, where most dramatic error happens when distance is low and *H* is also low. Introduction of *H* shows that contamination can result in both over and under-estimation of error. In particular, for larger values of *D*, if contaminants are similar between the two samples, relatively low levels of contamination can lead to sever under-estimation of distance. For example, with *D* = 0.18, if the samples are contaminated at 5% with somewhat similar species with *H* = 0.5, the estimated distance will be under-estimated by 43%.

**Figure 2:**
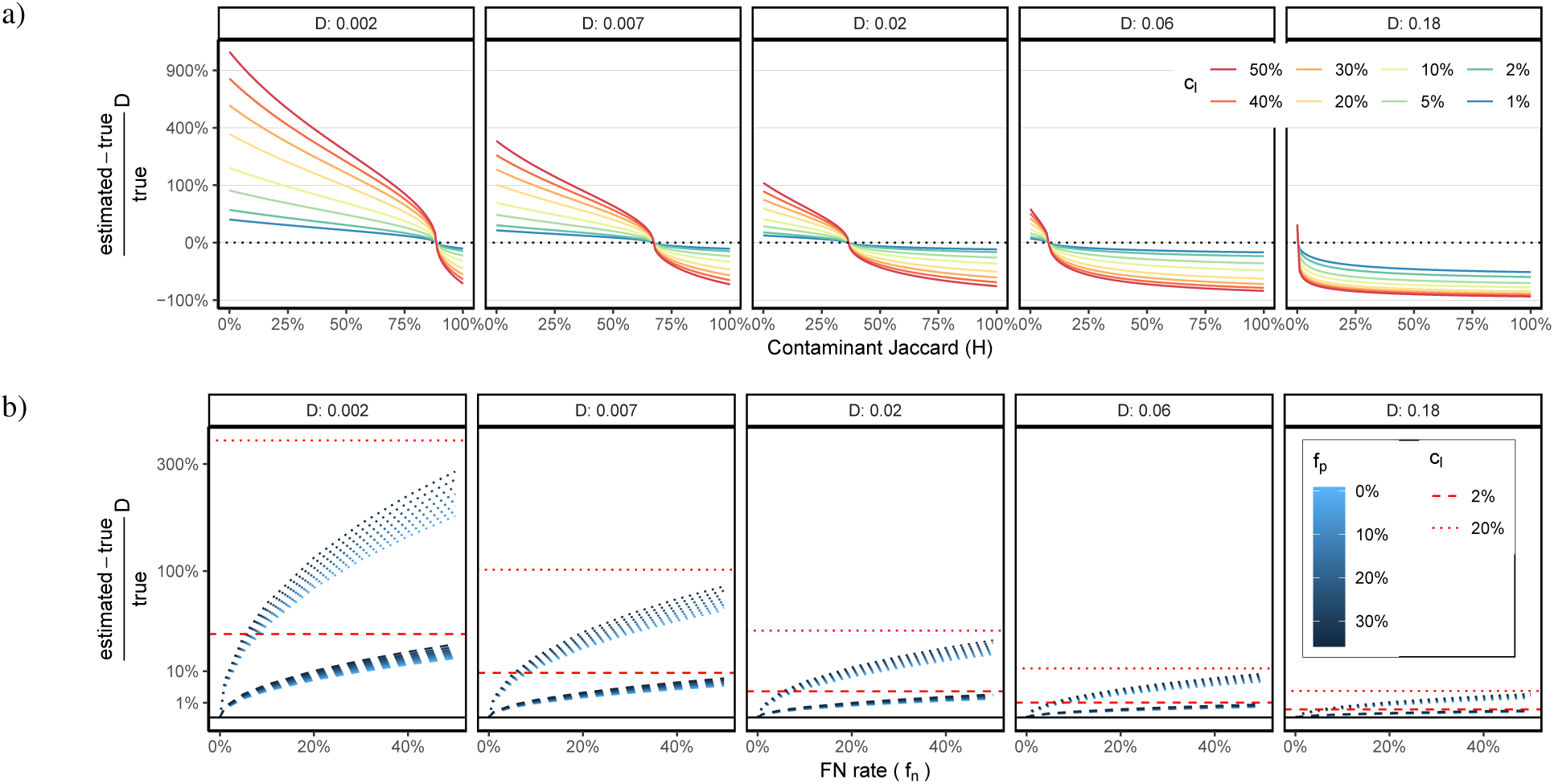
Theoretical modeling. (a) Impact of contamination on the genomic distance estimated from Jaccard according to theoretical expectation assuming contaminant *k*-mers of the two skims have a Jaccard of *H* (4). *For several D* and varying *H*, relative error is shown for eight contamination levels *c*_*l*_ = *c*_1_ = *c*_2_. (b) Error in Skmer distance (computed using (1), with Jaccard approximated using (5)) in the presence of filtering and with the disjoint contaminant *k*-mer assumption for various levels of FP portion (*f*_*p*_), FN (*f*_*n*_) rate, and *c*_*l*_. Red lines show the error in the absence of filtering. y-axis is in square root scale and *k* = 31.

#### 2.1.2 Impact of exclusion filtering

One approach to deal with contamination is using exclusion filters: search *all* reads in a genome skim against a (potentially incomplete) library of *known* contaminants and filter out reads that match the library. This approach will impose a trade-off between two types of possible errors. A false positive (**FP**) occurs when we incorrectly filter out a read that belongs to the target genome. A false negative (**FN**) occurs when we fail to filter out a read that belongs to contaminants, perhaps due to an insufficient similarity between the read and genomes included in the exclusion library. The exact choice of the method and parameters used for mapping reads to reference contaminant libraries, in addition to the composition of the reference library, create a trade-off between FP and FN error. The trade-off poses an important question: which type of error, FP or FN, is more damaging? Falling back on the disjoint contaminant *k*-mer assumption, we can approximate impact of FP and FN on *J* given one more assumption: A *k*-mer shared between the two genome skims is either kept or removed from both skims.

Let *f*_*p*_ be the portion of all *k*-mers that we remove by mistake (FP) and *f*_*n*_ be the portion of the contaminant *k*-mers that we fail to remove (FN). The proportion of *k*-mers shared between genome skims after filtering is (1 − *c*_*l*_)(1 − *f*_*p*_)(1 − *ρ*) (Fig. 1c). Additionally, (1 − *c*_*l*_)(1 − *f*_*p*_) + *c*_*l*_*f*_*n*_ of the *k*-mers in each set are retained after filtering for the total number of unique *k*-mers to be 2((1 − *c*_*l*_)(1 − *f*_*p*_)+ *c*_*l*_*f*_*n*_) −(1 − *c*_*l*_)(1 − *ρ*)(1 − *f*_*p*_). Thus,

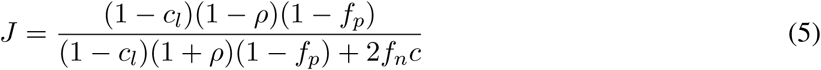

By plotting this equation as we vary the four parameters (*D, c*_*l*_, *f*_*p*_, and *f*_*n*_), we observe that filtering can successfully reduce the impact of contamination under many but not all conditions (Figs. 2b and S2). Filtering can be very effective in making Jaccard index close to what we would obtain without contamination, and overall, Jaccard is more sensitive to FN errors than it is to FP errors. Impact of filtering on genomic distance depends on the level of contamination, false negatives, and most of all, the true genomic distance. Reassuringly, in this model, the estimated accuracy of distance after filtering is reasonably high in most cases (Figs. S2 and S3). Nevertheless, in the most challenging cases, filtering cannot sufficiently reduce the error. With *D* = 0.002, unless *f*_*n*_ is low or *c*_*l*_ is moderate, error can be very high. Overall, *f*_*p*_ errors are less damaging than *f*_*n*_. Practically, it seems that with *f*_*n*_ ≤ 0.2, highly accurate estimates of distance are possible unless contamination levels are very high *and* the genomic distance is very low.

### 2.2 Empirical analyses using Kraken-II

Our empirical experiments validate the effectiveness of exclusion filtering, focusing on a leading *k*-mer-based read mapping tool called Kraken-II, originally designed for metagenomics and adopted here for contamination filtering. We start by describing Kraken-II and then detail the setup for the four experiments performed.

#### 2.2.1 Kraken

Kraken-II works by mapping all *k*-mers of a read to *k*-mers in a reference library and calls a read a match if the number of *k*-mers matching is *strictly* larger than a user-provided threshold called the confidence level (*α*). Kraken-II uses LCA-mapping to find lowest taxonomic level at which the read can be confidently matched. It also uses wild-carding *s* random positions of each *k*-mer (Brinda et al., 2015) to increase the sensitivity of matches. We will explore both *k* and *α* settings but fix other parametrs. We set minimizer length *l* = *k* or use the maximum allowed *l* value (31) for reference databases built with *k* > 31. We set *s*, the number of wild-card positions, to its maximum allowable value, *l*/4. We design our reference Kraken-II libraries to include a set of potential contaminants and as query, we use the bag of all reads in a genome skim (details described below). We will use microbial genomes to simulate contamination, and thus, all reference libraries we use are microbial. In contrast, our base genomes are Eukaryotic (plants or insects).

#### 2.2.2 Experiments

We present four experiments that explore the impact of *D* (equivalently, *ρ*), *c*_*l*_, *f*_*p*_, and *f*_*n*_. In addition, we test the running time of Kraken-II. Below, we describe the setup used in each experiment.

##### Exploring FN and FP of Kraken-II

We start by examining the sensitivity of Kraken-II to two parameters: *k* and *α*. We also consider completeness of the reference library, which is expected to have a direct effect on *f*_*p*_ and *f*_*n*_ rates. A lack of sufficiently close genomes to the contaminant can prevent Kraken-II from finding a match, and presence of genomes similar to non-contaminant genomes can cause FP matches. Thus, we define a third variable, *M*, as the genomic distance between a query and its closest match in the library. We control *M* by carefully selecting species included in the reference library and those used as query.

To control *M*, we use an available reference phylogeny of 10,575 bacterial and archaeal genomes (Zhu et al., 2019). Five genomes from this set had IDs that did not exist in NCBI anymore. We assigned remaining genomes to the reference library (10,460 genomes), the query set (100), or both (10). Based on the available phylogeny, we select 10 sets of 10 query genomes such that all genomes in a set have similar patristic (tree) distance to their closest leaf in the tree, not counting the query genomes. These sets had mean tree distance of {0.01, 0.02, 0.04, 0.06, 0.09, 0.10, 0.18, 0.23, 0.57, 1.20} and at most 25% divergence from the mean. We also randomly chose 10 genomes to be added to both reference and query sets. Then, for each of the 110 query genomes, we used Mash to compute *M*: its minimum distance to any of the 10,470 reference genomes. We then binned the 110 queries into 10 bins based on *M* (Table S1). Finally, we added 10 plant genomes (Table S9) to the set of query genomes in every bin. Plant species are from a different domain of life compared to the reference set and should not match the library; thus, they allow us to measure FP and TN rates.

We built Kraken-II reference libraries for selected *k* values (ranging from 23 to 35) using the 10,470 bacterial and archael reference genomes. Kraken-II only allows adding additional custom genomes of interest to its existing standard reference libraries. We used Kraken-II RefSeq viral genome database as a base library. All custom reference libraries were constructed without masking low complexity sequences.

We used the ART simulation tool (Huang et al., 2012) with HiSeq 2500 single read profile, 150bp read length with 10bp standard deviation to generate ≈1.4GB of synthetic reads (1000×coverage for each genome) for all query genomes. Every query genome was then downsampled to 1G for normalization purposes.

Reads in each query bin were queried against every constructed reference library for each *k* using several confidence levels (0 – 0.3). We then calculated TP, FP, FN, TN for every bin. TP is the count of bacterial/archael reads matched to Bacterial or Archaeal domains; FP is the count of plant reads matched to Bacterial or Archaeal domains; TN is the number of plant reads that are left unclassified by Kraken; FN is the number of unclassified bacterial/archaeal sequences. We use standard definitions of FPR=(FP)/(FP+TN) and Recall=TP/(TP+FN) and construct ROC curves in the standard fashion for every tested condition.

##### Skmer distances (simulation)

We next study the impact of contamination on distances computed from pairs of genome skims simulated from Drosophila assemblies. We first emulate the disjoint contaminant scenario by contaminating one of the two genome skims at a level *c*_*l*_. We used *D. simulans w501* to simulate the contaminated genome skim and used *D. simulans WXD1, D. sechellia*, or *D. yakuba* to simulate the uncontaminated skim. Based on assemblies, the distances between *D. simulans w501* and the three other species are 0.2%, 2.1%, and 6.3%, respectively, and we treat these as true distances. To add contamination, we use the same 110 query genomes described earlier but bin *M* into four ranges: [0, 0], (0, 0.05], (0.05, 0.15], and (0.15, 0.25], which include 10, 43, 19, and 17 species, respectively, corresponding to a total size of 37Mb, 76Mb, 37Mb, and 35Mb. Since our base Drosophila genomes are roughly 150Mb in size, we can add up to 25% contaminant reads for all bins, except for the (0, 0.05] bin, where we can add up to 60%. We concatenated all the genomes in each bin and used ART with the same settings indicated above to generate contaminant reads, which we then mixed with reads simulated from the main genome at levels varying from 0% to 60% (for the second bin) or to 25% (for all other bins) for a total of 0.1Gb per skim (thus, no more than 1× coverage). These read contamination levels translate to similar *k*-mer contamination levels (Table S3). We report the relative error in estimated distances as we increase the contamination level, both with and without Kraken-II filtering. Kraken is run with the same reference library used in the previous analysis.

We then simulate a scenario where *both* genome skims are contaminated with *overlapping* sets of species. Here, we only use the *M* ∈ (0, 0.05] bin and fix read contamination level to 15%. To control *H*, we randomly split bacterial reads into three parts: two unique parts and one part that served as an overlap. Every sample was generated by mixing unique and overlap contaminant portions with Drosophila genome skims at controlled ratios, with overlap set to 0% – 50%. Since unique parts can have evolutionary similar species, even the case of 0% overlap results in some *k*-mer overlap. Thus, we estimated contamination overlap (*H*) empirically using Jellyfish (Marçais and Kingsford, 2011) and saw it varied between 11% and 41% (Table S4). Finally, to have *H* = 0%, we added the disjoint set experiment with *M* ∈ (0, 0.05] and *c*_*l*_ = 15% to this set as well.

##### Skmer distances on real data

To move beyond simulations, we also evaluate effectiveness on real data with real contaminants. To do so, we utilized data from recent Drosophila assembly study by Miller et al., 2018. We subsampled available short read sequencing data (e.g., SRA files) to obtain 100Mb genome skims for 14 Drosophila species. We removed adapters, deduplicated and merged paired end reads using BBtools Bushnell et al. (2017). Then, we determined distances for all pairs of genomes before and after filtering them with Kraken-II. Distance error for every pair of genomes was estimated relative to the true distance defined to be the value computed by running Skmer on corresponding assemblies. In this experiment, we used a standard reference library available from Kraken-II distribution. This database includes RefSeq assemblies of all available bacterial, archaeal, viral and human (GRCh38) genomes as well as the UniVec_Core subset of the UniVec database (a total of 168483 genomes, as of July, 2019). We used default Kraken-II settings.

##### Impact of filtering on phylogeny

On the real Drosophila data, we also infer phylogenetic trees from distances and measure phylogenetic error. To estimate the phylogeny, we use FastME 2.0 software (Lefort et al., 2015) with JC69+Γ (Jin and Nei, 1990) model of evolution. Alpha parameter of JC69+Γ model is set equal to 1, which is the default value in FastME. We infer phylogenies from distance matrices obtained from assemblies and from genome skims before and after filtering. As the gold standard reference tree used for error calculations, we use the tree obtained from Open Tree of Life (OTL) (Hinchliff et al., 2015)(Fig. S4). We estimated branch lengths of the true tree using OTL tree topology and assembly distances under JC69+Γ model. We measure phylogenetic error using three metrics. (1) Normalized Robinson and Foulds, 1981 (RF) distance is total number of branches not matching. (2) Normalized weighted RF (wRF) distance is similar but each present or absent branch in each tree is weighted by the absolute difference between its lengths in the two trees, and then the total sum is normalized by the sum of branch lengths of the two trees. (3) Fitch-Margoliash (Fitch and Margoliash, 1967) is the weighted least squares error (FME) for species *i*, given as: *Q*(*i*) = Σ_*i*≠ *j*_ (*D*_*ij*_*/d*_*ij*_ − 1)^2^ where *D*_*ij*_ is the (corrected) distance between species *i* and *j*, and *d*_*ij*_ is sum of the branch lengths on the path connecting *i* and *j* on the phylogeny inferred using *D*. We also report cumulative FME of a phylogeny, which is 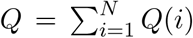. Denoting FME error on true and estimated phylogenies with *Q*(*i*) and 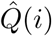 respectively, relative FME error is defined similarly to (3).

## 3 Results

### 3.1 Sensitivity of Kraken-II (FN and FP analysis)

The ability of the default version of Kraken-II (*k* = 35, *α* = 0) to find a match in the database is a direct function of *M*, the distance of the query to the closest match (Figs S5 and 3a). When the query has a close match in the library (e.g., *D <* 0.05), Kraken-II is able to match 80% – 100% of reads, which would result in tolerable *f*_*n*_ rates of 20% or less. As *M* increases, the ability of Kraken-II to classify degrades linearly with *M* up until around *M* ≈ 0.3 where Kraken-II fails to classify almost all reads (Fig. S5). Interestingly, when Kraken-II finds a match, it is often able to classify the read all the way down to the species level (Fig. S6).

Consistent with these results, when mixed plant/microbe skims are queried using the default Kraken, the recall of the filtering step is reasonably high (e.g., >85%) and the FP is low (4.5%) for *M* ≤ 0.05 (Fig. 3a). When 0.05 < *M* ≤ 0.1 or 0.1 < *M* ≤ 0.15, there is a substantial reduction in the recall to 67% and 56%, respectively; for 0.15 < *M*, recall is less than 33%, and thus filtering is not effective in those conditions.

**Figure 3:**
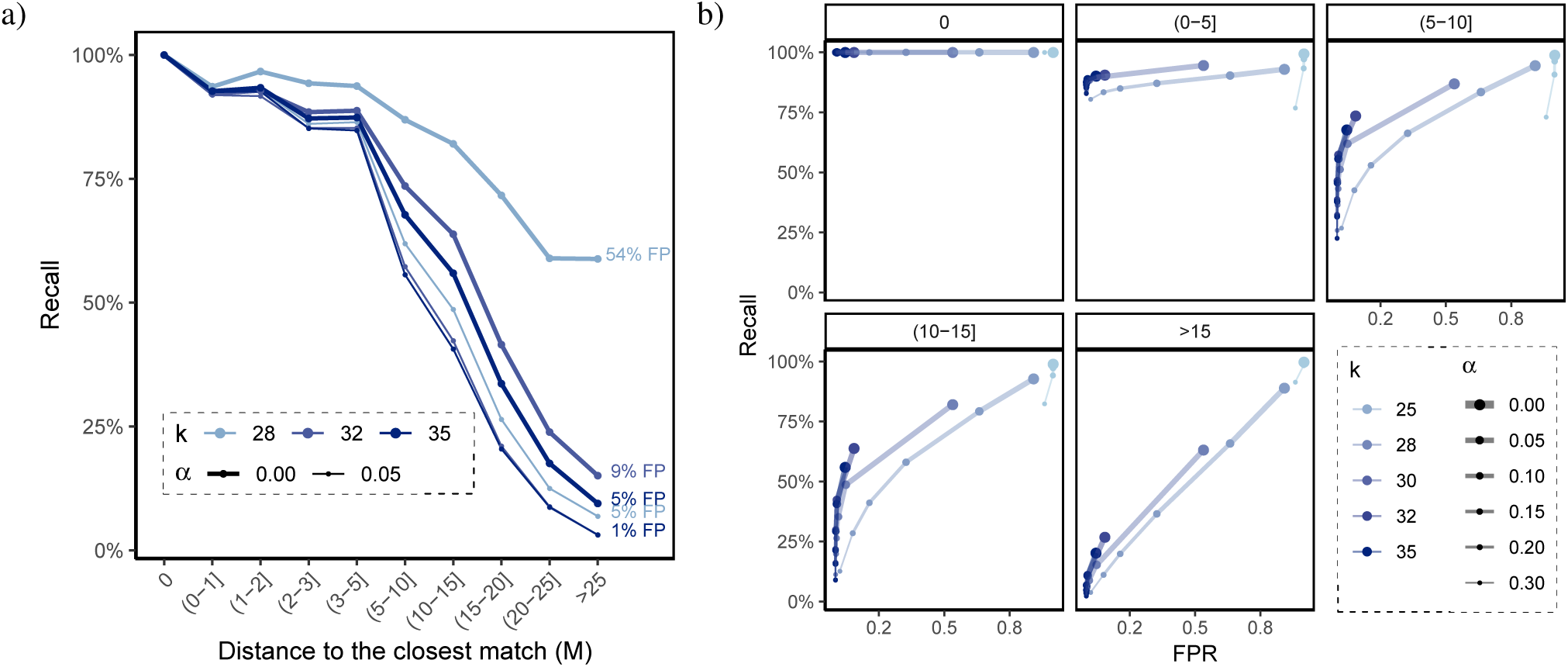
Sensitivity analysis of Kraken-II. (a) For three selected values of *k* and two selected values of *α*, the lines show recall for different bins of *M* (x-axis). Each line is labelled with its associated FPR. Note that FPR is a function of the plant genomes, which are identical across bins; thus, FPR is not a function of the match parameter *M.* (b) ROC curves for all *k* and *α* values across different bins of *M*, reduced to 5 ranges for ease of visualization (boxes). See Figure S7 for all 10 bins.

Given the low recall in some conditions and our expectation that FP error is less damaging than FN, one may aspire to increase the sensitivity of Kraken-II by adjusting its parameters *k* and *α*. However, our careful analysis of FP versus FN shows very limited ability to control the rates in a reasonable range (Figs. 3ab and S7). Many settings of *k* and *α* result in FP error above 50% and often close to 100%. Many of the settings also have high FP without improving recall compared to default settings (Figs. 3b). The only settings that seem to provide a reasonable trade-off between FPR and recall are *k* ∈ {35, 32, 28} and *α* ≤ 0.05. Focusing on these settings (Fig. 3a), we observe that setting *k* = 28 and *α* = 0 provides a substantial increase in recall but increases FPR to unacceptable levels (55%). *k* = 32 improves recall compared to the default setting for *M* ≥ 0.05 bins by a consistent but relatively small margins (5–8%), but also increases the FPR to 8.5%. Overall, changing parameters do not result in substantial improvements over the default settings.

### 3.2 Impact of filtering on Skmer distances (simulated contaminants)

#### Disjoint contaminants

Focusing on simulated contamination between pairs of Drosophila genome skims, when only one species is contaminated, increasing the contamination level results in increasing error in estimated Skmer distances, going up to 90% error for *D* = 2% and 1000% for *D* = 0.2% when *c*_*l*_ = 60% (Fig. 4a). As theory suggested, here, the strongest detrimental effect appears for *D* = 0.2%.

**Figure 4:**
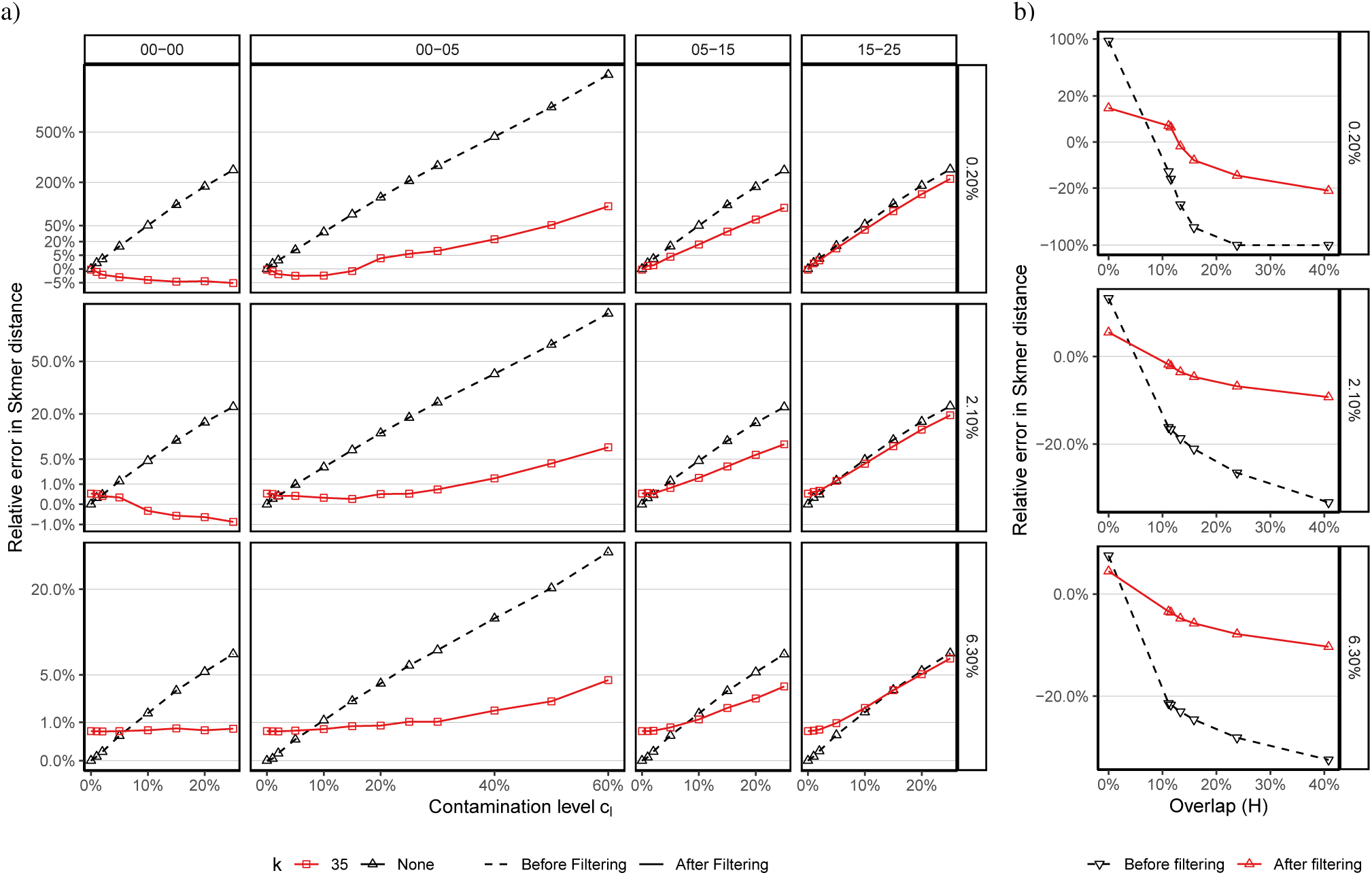
Filtering on simulated Drosophila genome skims. Relative error of Skmer distances without (dashed) and with (solid) Kraken-II filtering. Three pairs of Drosophila are chosen to be at true distance *D* = 0.2%, *D* = 2.1%, or *D* = 6.3% (rows). Contaminants are selected such that they are at distance *M* from their closest match in the reference contaminant library. (a) Simulating contaminants in only one of the two species (disjoint contaminants) for four ranges of *M* (columns) and various levels of contamination (x-axis). (b) Contaminating both genomes such that the overlap between contaminants measured by Jaccard similarity is 0% ≤ *H* ≤ 41%. Here, *c*_*l*_ = 15% per species and *M* ∈ (0, 0.05]. Y-axis is on square root scale; see S8 for normal scale and a range of *k* and *α* values.

Filtering using default Kraken-II dramatically reduces the error *when* the contaminant has an exact or close match in the reference library (Fig. 4a). For *M* ≤ 0.05, remarkably high levels of contamination are tolerated after filtering. For example, for 0 < *M* ≤ 0.05 and *D* = 2.1%, even with high *c*_*l*_ in 25% – 50%, distances have only 0.3% – 4% relative error after filtering. For *D* = 6.3%, error after filtering is never more than 5% for *M* ≤ 0.05. Even in the most challenging case of *D* = 0.2%, *c*_*l*_ = 25% leads to only 6% error after filtering in contrast to 206% error before filtering. Despite the improved accuracy overall, in some cases, filtering can increase the error slightly but noticeably, perhaps due to FP filtering of correct reads. For *M* = 0 and *D* = 6.3%, if contamination is below 5%, no filtering is better than filtering, which always results in ≈0.6% relative error regardless of the level of contamination. Interestingly, in some cases, filtering can result in *underestimation* of distances (e.g., up to 1% for *D* = 2.1% and *M* = 0).

In contrast, for contaminants without a close match with *M >* 0.05, filtering fails to fully remove contaminants. Nevertheless, for 0.5 < *M* ≤ 0.15, filtering has substantial benefits. For example, error is reduced from 180% and 16% with no filtering to 65% and 6%, respectively for *D* = 0.2% and *D* = 2.1%. These reductions, while substantial, may not be sufficient. Even worse, for *M >* 0.15, filtering has very little or no ability to reduce the error and decreases or increases the error by very small margins.

Finally, changing *k, α* settings of Kraken-II does not consistently improve the accuracy above and beyond the default setting (Fig. S8). Using *α* = 0.05 can very slightly reduce the error for the *D* = 6.3% case but is not dramatically different. Thus, we will exclusively use the defaults in the next experiments.

#### Overlapping contaminants

When both skims are contaminated with overlapping species, as theory suggested, we see *under-estimation* of distances (Fig. 4b). These under-estimations can be dramatic, going all the way down to −100% (i.e., the estimated distance is 0). Once again, filtering using Kraken is able to improve results dramatically, resulting in relative error that does not exceed 23% for *D* = 0.2% and is at most 6% in the remaining cases.

### 3.3 Impact of filtering on Skmer distances (real contaminants)

In the experiment on real unassembled Drosophila sequences, absent any filtering, Skmer often under-estimates distances (Fig. 5a). The under-estimation of distances is consistent with our theory assuming *H >* 0 (Fig. 2). Kraken-II run on these data identifies between 5.5% and 15.1% of the reads as belonging to human or microbes (Fig. 5b). Interestingly, for most Drosophila species, Kraken-II assigns ∼40–50% of the matched reads to one of three genera (Homo, Acetobacter, and Clostridium), indicating that many pairs of genome skims have similar contaminants (i.e., *H >* 0). Therefore, the under-estimations of distances matches the theory. Consistent with this explanation, we observe that the error in computed distances is associated with the percentage of the reads found by Kraken-II to be of human or microbial origin (Fig. S9).

**Figure 5:**
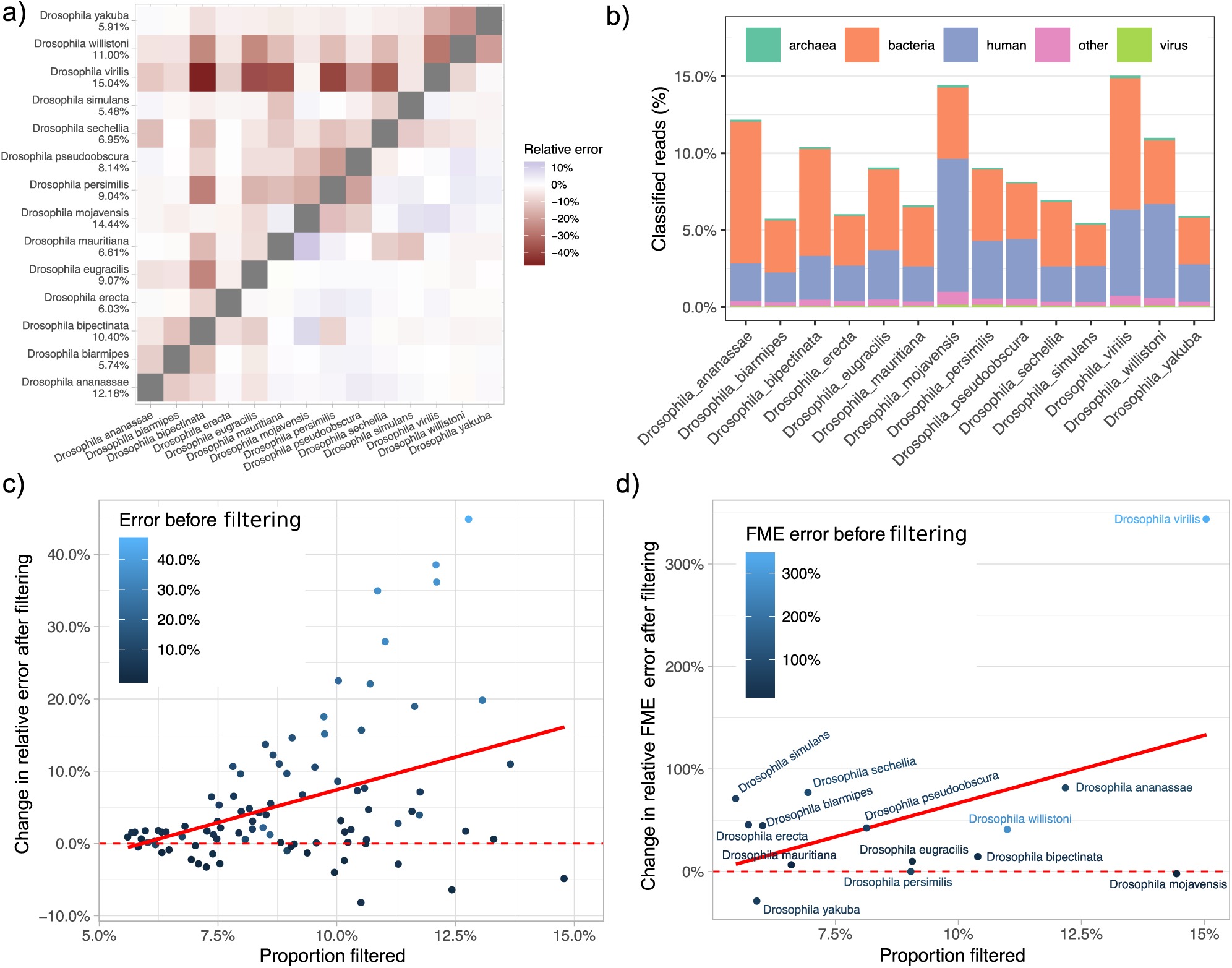
Filtering of real contaminants. (a) Relative distance error before (upper triangle) and after (lower triangle) filtering per pair of Drosophila species. Numeric lables on y-axis represent percentage of reads filtered per species. (b) Percent of reads classified by Kraken-II to different groups. “Other” corresponds “cellular organisms” (shared between domains). (c) Change in the relative distance error after filtering. Positive values indicate a reduction in error. (d) Change in relative FME error per species after filtering. Solid red line: a trend line fitted to the points.

Filtering reads using Kraken-II dramatically reduces the errors in Skmer distances (Fig. 5c). Over all pairs, the mean absolute relative error reduce from 9.1% before filtering to 3.4% after filtering. In some cases, reductions are dramatic. For example, the relative error in pairwise distances between *D. virilis* and *D. bipectinata, D. eugracilis* and *D. mauritiana*, decreased from 46.2%, 36.9% and 35.9% before filtering to 1.3%, 0.8%, and 1.0% after filtering. In a minority of cases, error increased after filtering but the increase in error never exceeded 8% (*D. mauritiana* vs. *D. mojavensis*) while reductions in error could be as high as 45% (*D. virilis* vs. *D. bipectinata*) (Fig. 5c). The wide range of error reductions is unsurprising given that the actual level of contamination in original sample can vary substantially. The magnitude of improvement in distance estimates is positively correlated with the percentage of reads filtered (Fig. 5c), the genomic distance (Fig. S10a), and the magnitude of the error before filtering (Fig. S10b).

When there is error after Kraken-II filtering, it tends to be due to *over-*estimation of distance, as opposed to under-estimation observed before filtering, suggesting that Kraken-II perhaps over-filters reads. Most extreme cases of over-filtering involve distance estimates between a single species, *D. mojavensis*, and other species such as *D. bipectinata, D. mauritiana* and *D. virilis*. The *D. mojavensis* is the only species with high levels of Kraken-II filtering but low error rates in pairwise comparisons. Interestingly, *D. mojavensis* also includes the highest levels of contamination from unknown sources.

### 3.4 Impact on phylogenetic reconstruction

The phylogeny inferred from Skmer distances computed from the assembly and modeled using JC69+Γ is topologically identical to the gold standard OTL phylogeny, and its total FME error is only 0.03 (Fig. S4). However, the phylogeny estimated using the same method but using genome skims has two wrong branches (RF = 4), and a FME of 1.26. Thus, absent filtering, genome skims produce trees with substantial error.

Improvements in estimated genomic distances due to filtering translate to improved phylogenetic trees. The tree topology improves only slightly and has one incorrect branch (RF = 2) after filtering. However, the improvements in estimated branch lengths, as reflected in wRF and total FME error, are dramatic. Filtering leads to nearly 70% decrease in total FME metric, from 1.26 to 0.38, and a similar level of reduction is observed for wRF (Table S5). Examining individual branch lengths, the phylogeny using filtered data is much more similar to the true tree (Fig. S4).

When we use FME to measure the impact of filtering on the phylogenetic error of individual species, we observe patterns consistent with reductions in distance error (Fig. 5d). Individually, majority of species have reduced FME after filtering, with the most extreme FME reduction happening for *D. virilis* by nearly %350. Consistent with previous results, we observe that the FME error of *D. mojavensis* does not decrease (but it also does not increase). As expected, gains in phylogenetic error measured by FME correlate with the amount of filtering performed by Kraken (Fig. 5d).

### 3.5 Running time

We assessed running time performance of Kraken-II on skims of five randomly selected species from different domains of life (Table S6) using both single-threaded and multi-threaded (24 threads) modes of operation. Run time was found to linearly increase with the skim size, regardless of the number of threads used (Fig. S11). With 1Gb of reads, the running time of the single-threaded version was below 100 seconds on an Intel Xeon CPU in all cases we tested. Running Kraken-II with 24 threads reduced the speed by a factor of 10, offering a significant improvement. The main limitation with Kraken-II is its significant memory requirement during queries, which requires between 100Gb and 120Gb for our reference libraries.

## 4 Discussion

The use of genome skimming in the literature has mostly relied on assembled organelle genomes (e.g., Coissac et al., 2016; Dodsworth, 2015; Malé et al., 2014; Weitemier et al., 2014). These approaches rely on assembly construction pipelines (e.g., Jin et al., 2019) to remove contaminants (i.e., to avoid mis-assembly or to filter out mis-assembled contigs). Elsewhere, we have advocated going beyond organelle genomes and using all reads in an assembly-free fashion to increase the resolution of taxonomic identification (Balaban et al., 2019; Sarmashghi et al., 2019). However, this goal has been hampered by the presence of contaminants. This study showed a relatively effective way of dealing with contamination, hence bringing genome skimming based on nuclar reads one step closer to a reality.

Our study showed that Kraken-II is able to find contaminants that are within 5–10% genomic distance to the closest match in a reference library in a computationally efficient manner. Our modeling showed that FP errors were perhaps less detrimental to distance calculations than FN. Analysis of different *k* and *α* parameters did not reveal parameter combinations that could improve upon the using default settings of Kraken-II (*k* = 35, *α* = 0). Analyses of real data demonstrated that contamination removal can dramatically improve Skmer distance estimates in the presence of contaminants. These more accurate distance metrics computed after filtering can lead to reduced phylogenetic branch length error by up to 70% and can also improve the tree topology.

### 4.1 Usefulness of Theoretical Models

Simplified assumptions allowed us to establish theoretical models of the impact of contamination on estimated distance. The theory predicted that even small levels of similarity between contaminants (*H*) can lead to substantial under-estimation of distance when distance is large. Consistently, on the real data, where distances are often *>* 0.10, we observe under-estimation by 5% or more in 47 out of 91 pairs. Our results also showed high levels of similarity between contaminants of Drosophila genomes (where three genera made up 40–50% of contaminants). Thus, there is a reassuring match between the theoretical model and the observed data.

Kraken filtering improved accuracy on simulated and real data. On real data, it occasionally over-corrected errors, leading to over-estimation of the distance. These may be due to FP filtering, reduced coverage after filtering, or other factors not fully understood here. In our runs of Kraken-II on real data, we observed 5%–15% filtering. The lower value can be explained by ∼5% Kraken-II FP rate when run under its default setting. The upper value is consistent with ∼10% contamination level, a scenario that can easily happen in real sequencing projects.

Another potential use of the theory could have been developing filter-free methods of dealing with contamination. Just as impact of coverage and error on Jaccard can be modelled, we can compute the Jaccard index with no filtering but *correct* for the modelled impact of contamination on Jaccard. Given reliable estimates of *H, c*_1_, and *c*_2_, we can manipulate (4) to update (1) and obtain:

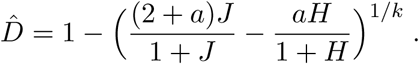

Adding coverage and error models, we can update (2) to:

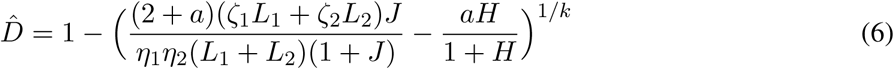

This equation allows for filter-free contamination-aware distance calculation. Unfortunately, however, this equation is extremely sensitive to correct estimation of all parameters, including *H, c*_1_, and *c*_2_ (Fig. S12). Even small mistakes (1-5% relative error) in the estimated contamination level or Jaccard can lead to dramatic errors in the estimated distance computed using (6). Since computation of these parameters is noisy, we do not advocate this filter-free method despite its theoretical elegance.

### 4.2 Filtering methods

Filtering requires a tool to answer queries of the following type: “Does this particular read belong to one of the genomes in a given reference library?”. We chose Kraken-II for answering these queries because of its high accuracy and reasonable scalability, as established in several bench-marking studies from the metagenomics field (McIntyre et al., 2017; Meyer et al., 2019; Sczyrba et al., 2017; Ye et al., 2019). It is also one of the most widely-used tools, with an active user support and stable and robust software development. Further, we explored three parameters of Kraken-II: *k*-mer length, confidence score and database content. Other parameters such as minimizer length (mostly relevant to storage and not accuracy) and minimizer space count are not explored here (Brinda et al., 2015). In our experiments we kept the number of wild-carding positions at its recommended upper limit and turned masking off but there might be a set of settings which in combination with masking can produce a more optimal sensitivity. We note that results from Wood et al., 2019 have indicated that Kraken-II is not very sensitive to particular parameter settings.

Alternatives to Kraken-II exist, and future studies can compare them to Kraken-II for genome skimming. BLAST (Altschul et al., 1990) and MegaBLAST (Morgulis et al., 2008) are the obvious alternatives but are an overkill for our problem. These tools perform alignment and can yield higher sensitivity than Kraken-II but are orders of magnitude slower (Wood and Salzberg, 2014; Ye et al., 2019). However, they produce more precise results (maps to individual species) than what we need.

Beyond alignment tools, most alternatives to Kraken-II are also *k*-mer-based, but differ in the way reference library is constructed and how the query is run. *k*-mer-based methods inculde LMAT (Ames et al., 2013), and CLARK(-S) (Ounit and Lonardi, 2016; Ounit et al., 2015). Benchmarking studies (e.g., McIntyre et al., 2017; Meyer et al., 2019; Sczyrba et al., 2017; Ye et al., 2019) do not indicate any consistent advantage in using these methods over Kraken-II, and many of them are slower. Among sufficiently fast tools are KrakenUniq (Breitwieser et al., 2018), Braken (Lu et al., 2017), and Centrifuge (Kim et al., 2016). KrakenUniq is recommended for use in cases where FP can be detrimental (e.g. in pathogen identification/diagnoses), but our theory and empirical data suggest FP is less important and FN in our application. Braken (Lu et al., 2017), an extension of Kraken-II, is focused on improving aggregated abundance profiles, a feature that is irrelevant to our usage. Centrifuge (Kim et al., 2016) uses FM-index lookups and within-species compression for mapping a read to one or more species. Compared to Kraken-II, Centrifuge is slower and needs more time for building its reference database. We leave its comparison to Kraken for future work.

A separate set of *k*-mer-based methods have been developed for finding RNAseq experiments that include a specific *k*-mer. Solomon and Kingsford (2016) introduced Sequence Bloom Tree (SBT) to allow very fast queries of a *k*-mer versus a reference set of experiments by creating a hierarchy of compressed bloom filters that store *k*-mers. Mantis (Pandey et al., 2018) is an alternative to Bloom filters based on counting quotient filters and is reported to be more memory efficient and faster that SBT-based methods. While these tools have been developed mainly for RNASeq analyses, in the future, they can perhaps be adopted for mapping reads to genomes with minimal changes to the algorithm. In fact, Kraken might implement counting quotient filter data structure in its future releases (Wood et al., 2019).

Beyond these tools, many other metagenomic methods have been designed for finding the taxonomic composition of a mixed sample (e.g., Liu et al., 2010; Milanese et al., 2019; Nguyen et al., 2014; Segata et al., 2012). However, these tools do not seek to classify *every* read from anywhere in the genome; they are either marker-based or use composition data. Thus, these tools are irrelevant to our queries.

### 4.3 Remaining gaps

In our study, we focused solely on prokaryotic and human contamination. Real contamination is more complex and can include eukaryotic microorganisms, traces of endosymbionts and diet, and various forms of lab contamination. Thus, many applications will benefit from more inclusive Kraken-II contaminant libraries. At a minimum, fungi need to be considered, especially for plants. Moreover, removing reads from organelle genomes, which are expected to be over-represented, may further improve accuracy.

Luckily, Kraken-II enables a straightforward mechanism for extending reference libraries. Our future efforts will include building a larger library of potential contaminants that includes fungi and perhaps expected sources of diet. However, such libraries will have to be group specific; for example, for skimming insects, we can treat plants as contaminants whereas in skimming plants, we should treat insects as contaminants. Ideally, individual genome skimming reference libraries for a target group (e.g., all insects) should be furnished with a relevant contaminant library especially designed for that group based on the knowledge of taxonomic groups expected to be present in its diet and its endosymbiont. Clearly, this approach runs into its limitations when endosymbionts or the diet happen to be from species with similar genomes to the target species.

The fundamental limitation of our exclusion filtering approach is that we need to know what broad group of species is expected to contaminate. This limitation is a result of our implicit assumption that a read is correct unless we find evidence to the contrary. Even when such biological knowledge is available – it may not be – this approach can fail to capture lab-introduced contamination (e.g., a plant species that was contaminated with fish due to failures in sample preparation or sequencing on the same lane).

Inclusion filtering is an attractive alternative to exclusion filters. Given a reference database of purified (perhaps using exclusion filters) genome skims, we can build a Kraken reference library from species in the skimming reference library. Then, for every new query genome skim, we can use that library to find reads that seem to match the broad taxonomic group of interest and only *include* those reads in the calculation of Jaccard. Our results indicate that this method would work only if the skimming reference database is so dense that each new query skim is expected to have a close match (e.g., < 5%) to one of the reference skims. Moreover, this approach is predicated on the reference library being free of contaminants. Despite these shortcomings, we believe this approach should be further explored in the future.

Finally, better algorithms for read matching seem necessary. Our results showed that Kraken-II provides a reasonable solution. Nevertheless, the method remains incapable of finding domain level matches when the closest match is moderately distant from the query. We believe it is possible to design more sensitive read mapping techniques that can match a species even when its closest match is *>* 10% distance. Note that in genome skimming, we are only interested to know whether a read belongs to a large taxonomic group, as opposed to metagenomics, when abundances and exact matches are desired. Given the less demanding needs of the skimming application, we anticipate that better algorithms can be developed in future to increase recall with little or no loss of specificity and speed.

## Availability of data and materials

Scripts and summary data tables are publicly available on https://github.com/noraracht/kraken_scripts.git. Raw data used in the manuscript is deposited in https://github.com/noraracht/kraken_raw_data.git. The detailed description of genomic datasets used in our experiments, accession numbers of the assemblies and the exact commands used to simulate genome skims are provided in Supplemental Material.

## Author Contributions

All authors conceived the idea. SM and VB developed a theoretical model. ER implemented the pipeline and performed experiments. MB completed phylogenetic experiments. All authors contributed to the analyses of data and the writing. All authors read and approved the final manuscript.

## Appendix A Derivations

### Appendix A.1 Derivation of (4)

#### Definitions

- Let *L* be the number of unique *k*-mers in the base genomes of species of interest, which for each of calculation, we assume is the same between genomes.
- *ρ* is the fraction of L *k*-mers that are different between the two base genomes.
- *Y*_1_ and *Y*_2_ are the total number of unique *k*-mers from each of the two contaminants.
- *c*_1_, *c*_2_ are the fraction of *k*-mers in sample one and sample two that come from contaminants. Thus, 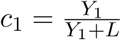 and 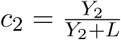.
- 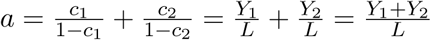. Thus, *Y*_1_ + *Y*_2_ = *aL*.
- *X* is the total number of *k*-mers shared between the two contaminant sets, and *H* is the Jaccard similarity between *k*-mers of the contaminants of the two skims. Thus,

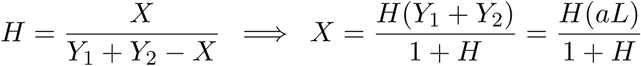

#### Goal

We want to derive

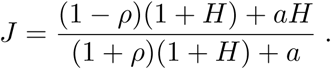

#### Derivation

We assume coverage is high enough that all *L k*-mers from each genome are in the skim. Then,

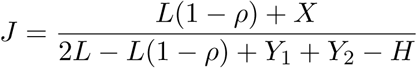

where the numerator counts the number of shared *k*-mers from the two base genomes plus the number of shared *k*-mers from the contaminants, and the denominator counts the total number of *k*-mers. Results are obtained as follows.

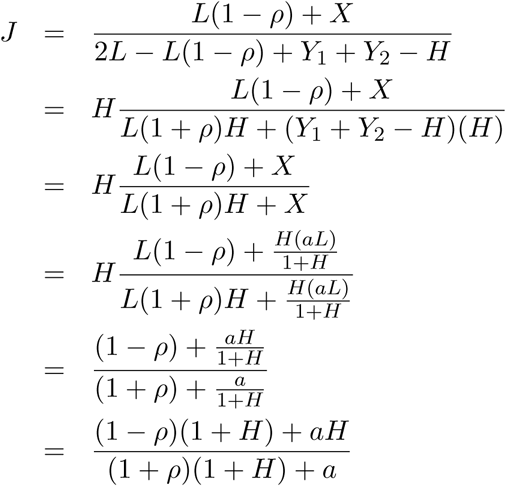

## Appendix B Supplementary Figures

**Figure S1:**
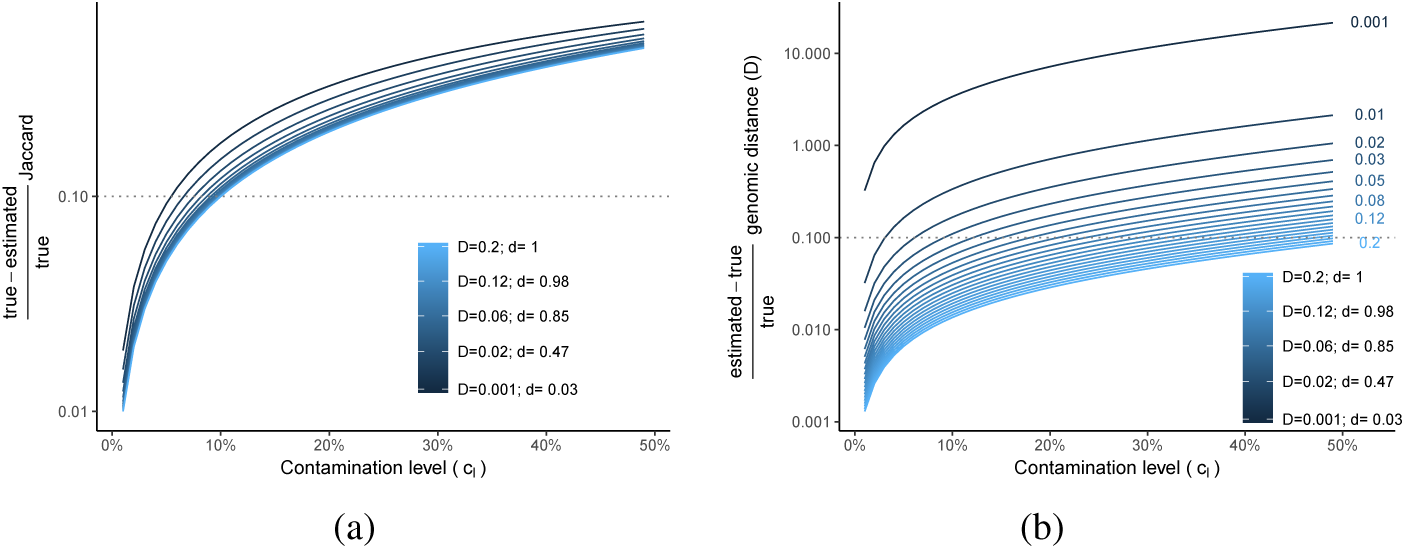
Theoretical modeling. Impact of contamination on Jaccard (a) and the genomic distance estimated from Jaccard (b) according to theoretical expectation under the disjoint contaminant *k*-mer assumption. For various genomic distance (*D* ∈ {0.001} ∪ {0.01, 0.02, …, 0.2}) corresponding to 0.03 < *ρ* < 0.99 and contamination levels 0.01 ≤ *c*_*l*_ ≤ 0.5, the relative error of the Jaccard index (a) and the estimated Skmer distance (b) as a result of ignoring contamination are shown. *k* = 31.

**Figure S2:**
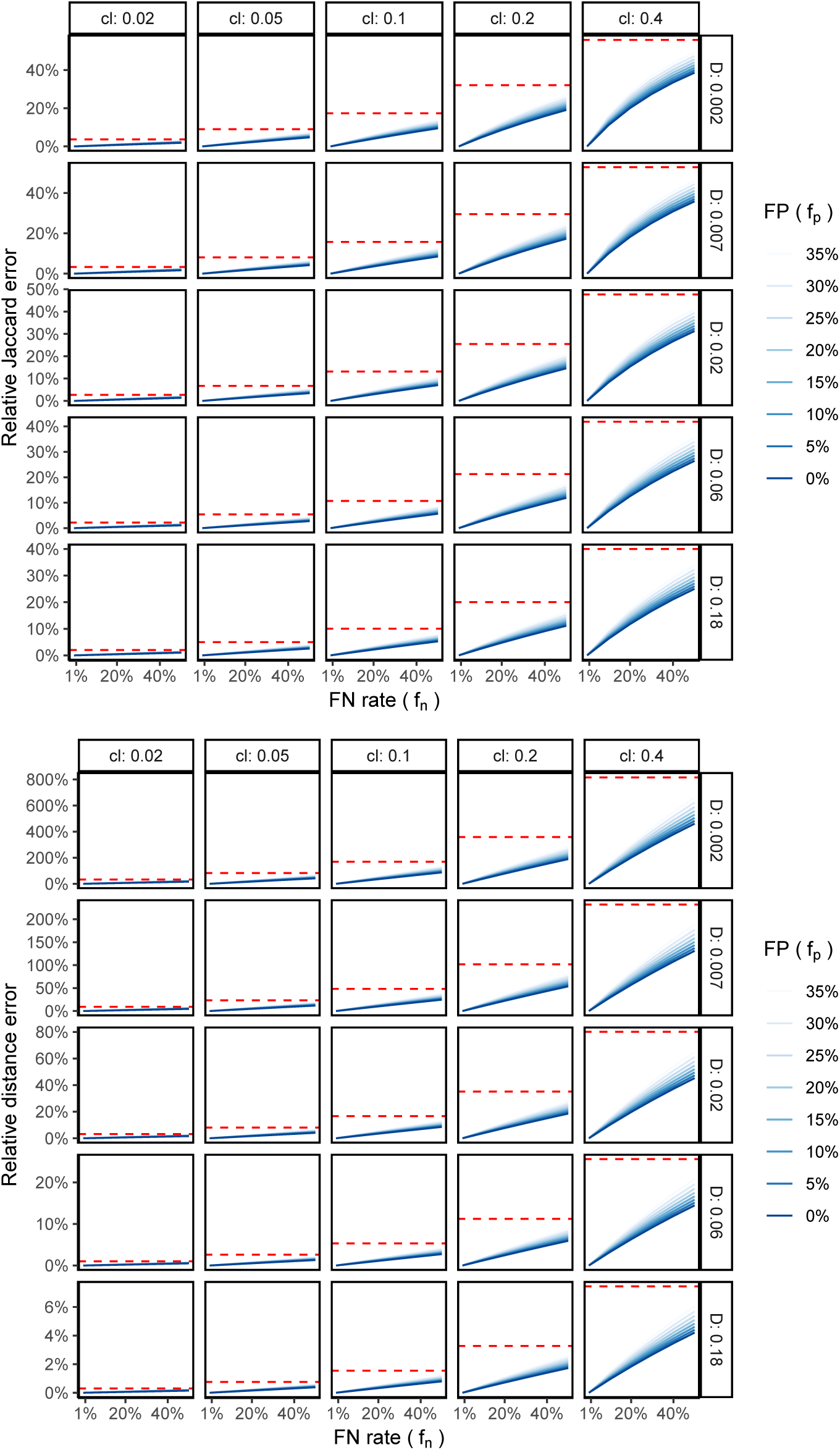
Theoretical impact of filtering on Jaccard (top) and Skmer distance (bottom). Two genomes with *D* between 0.001 and 0.05 are both contaminated at 0.01 ≤ *c*_*l*_ ≤ 0.32. Skmer distances are approximated using (1), with Jaccard approximated using (5). Results are for various levels of FP portion (*f*_*p*_), and FN (*f*_*n*_) rate. Solid lines show the relative error in Skmer distance after filtering, normalized by the true uncontaminated value, expressed as percentage. The error in the absence of filtering is shown as a horizontal dashed red line. See Figure S3 for a tabular view.

**Figure S3:**
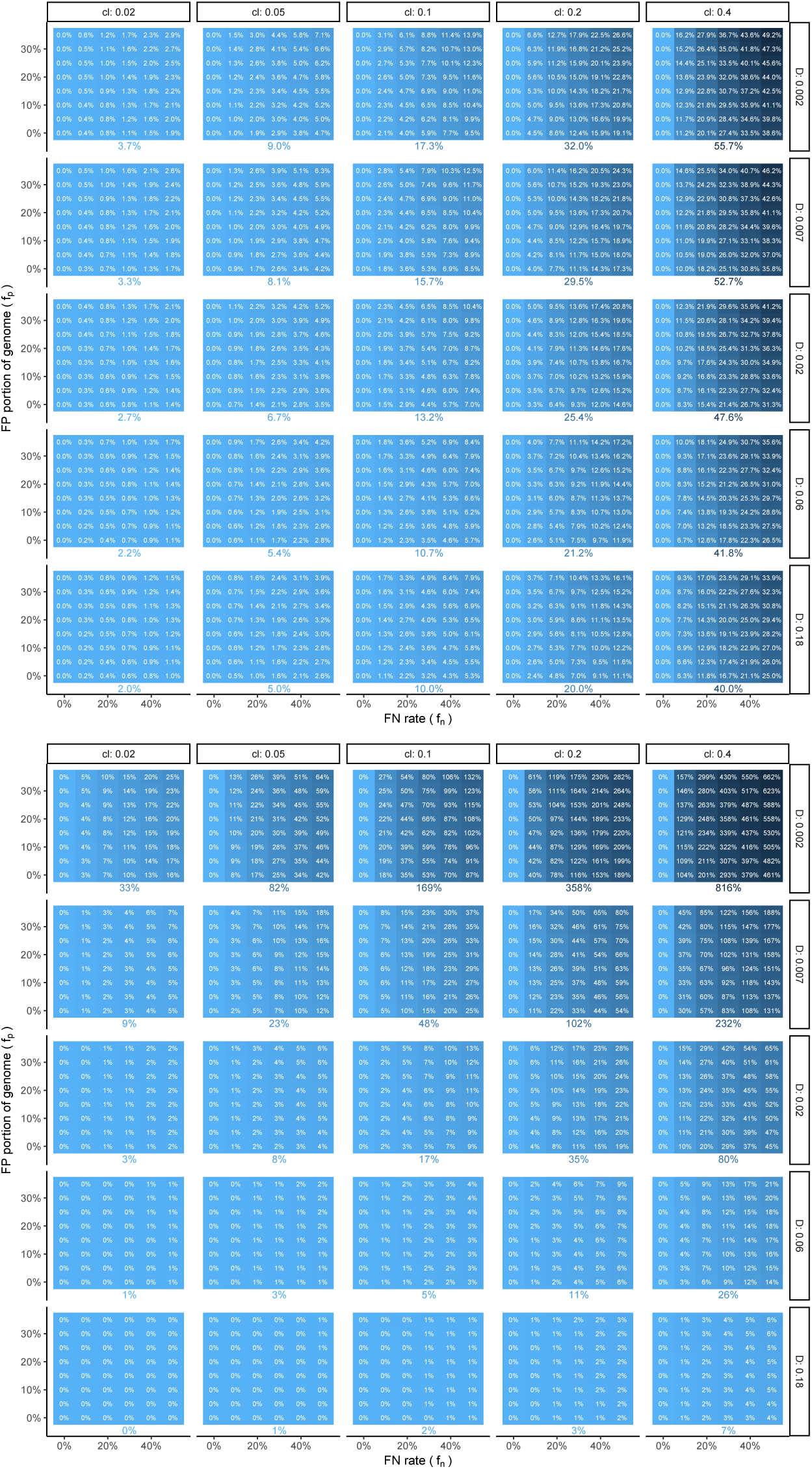
Approximate impact on Jaccard (top) and distance (bottom). Two genomes with portion 0.05 ≤ *ρ* ≤ 0.75 of their *k*-mers not matching (corresponding to *D* between 0.002 and 0.044) are both contaminated at 0.02 ≤ *c*_*l*_ ≤ 0.32. Approximation of Jaccard using (5) Skmer distance *D* using (1) for various levels of FP portion (*f*_*p*_), defined as the percentage of each genome skim that is filtered out by mistake, and FN Rate (*f*_*n*_), defined as the proportion of the contaminating *k*-mers that have *not* been removed. Each box shows the error in Jaccard or distance estimation after filtering, normalized by the true value (i.e., value with no contamination), expressed as percentage. The error in the absence of filtering is shown as a single number below each box.

**Figure S4:**
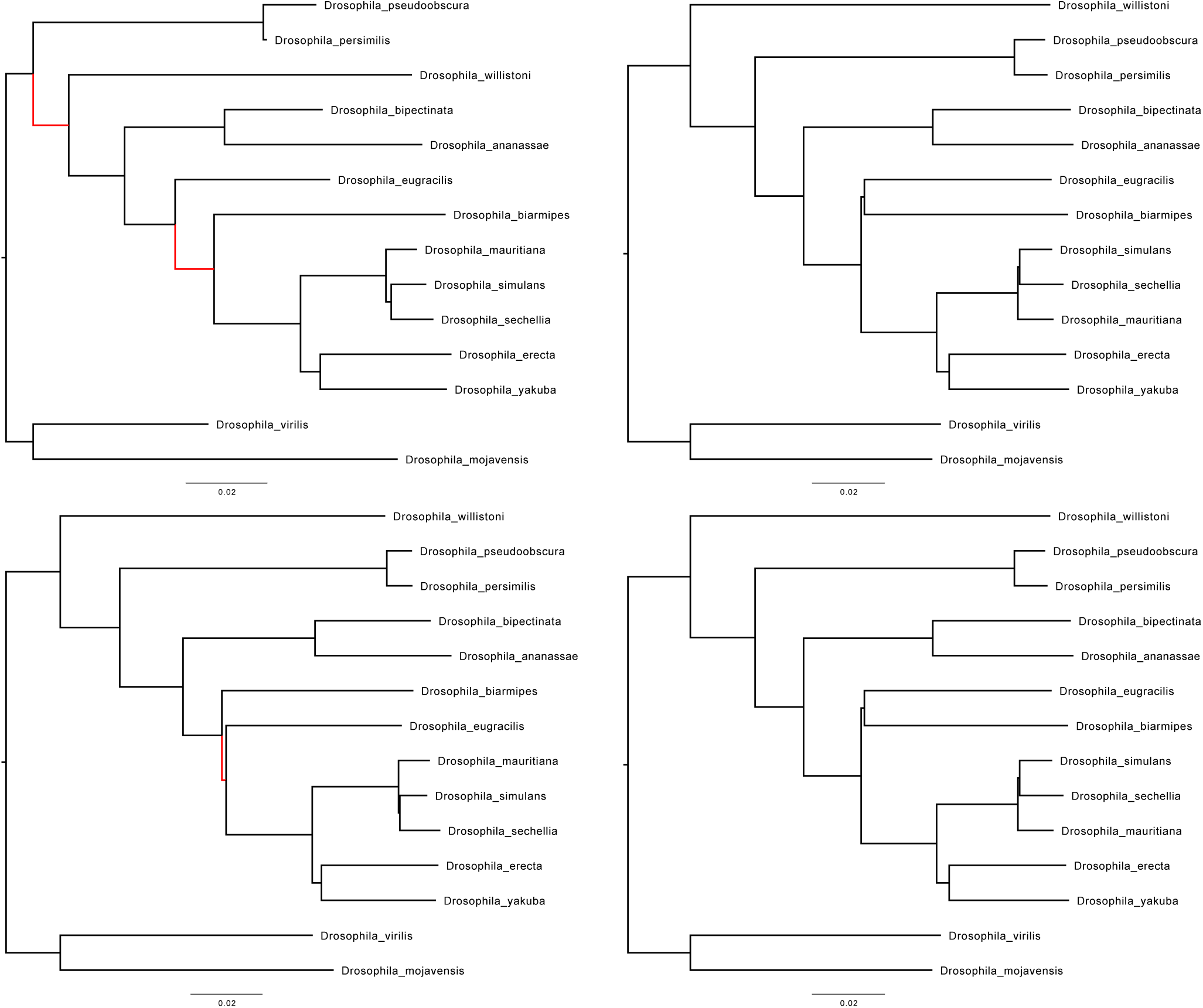
Drosophila phylogeny inferred from distances computed without filtering (top, left), and after filtering with Kraken-II (bottom, left) on 100Mb genome skims. Gold standard Drosophila phylogeny obtained from Open Tree of Life, whose branch lengths are computed using assembly distances, is show twice (top and bottom, right). All trees are based on Jukes-Cantor model of evolution accounting for rate variation across sites using Γ model with *α* = 1 and are inferred using FastME. Branches that do not match the gold standard phylogeny are indicated with red.

**Figure S5:**
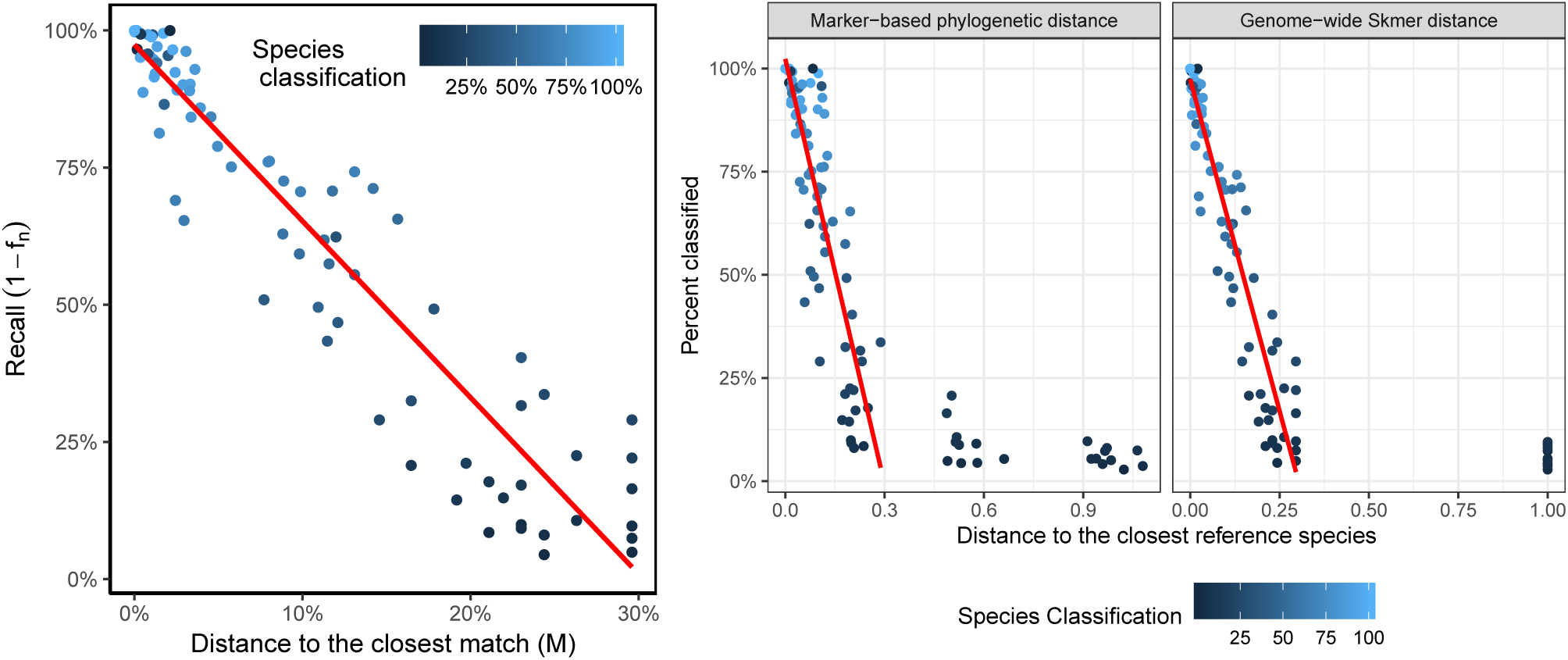
Sensitivity analysis of Kraken-II. (a) For each query genome, dots show the percentage of reads classified (i.e., 1 − *f*_*n*_) by the default Kraken-II at the domain level or lower versus the distance of a query to the best match in the reference library (*M*), measured using Skmer. Default Skmer is not accurate for *M >* 0.3 and thus we show *M* ≤ 0.3. (b) Similar to part (a), except, here, on the left, we measure *M* using either a phylogeny inferred from 381 marker genes and applying the inverse of the JC69 correction or by applying Skmer to the base assemblies. Using phylogenetic distances allows us to measure *M >* 0.3.

**Figure S6:**
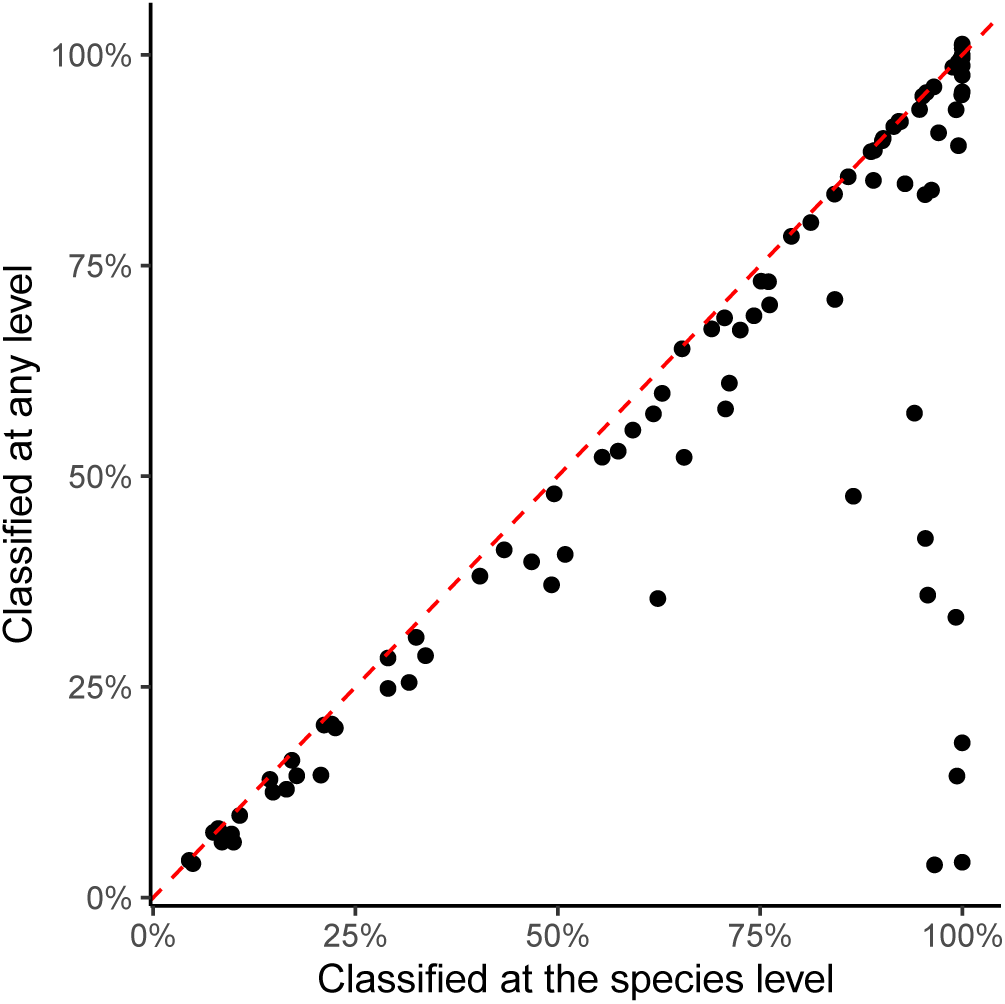
Sensitivity analysis of Kraken-II. For each query genome, dots show the percentage of reads classified by Kraken-II vs percentage of reads classified at the species level.

**Figure S7:**
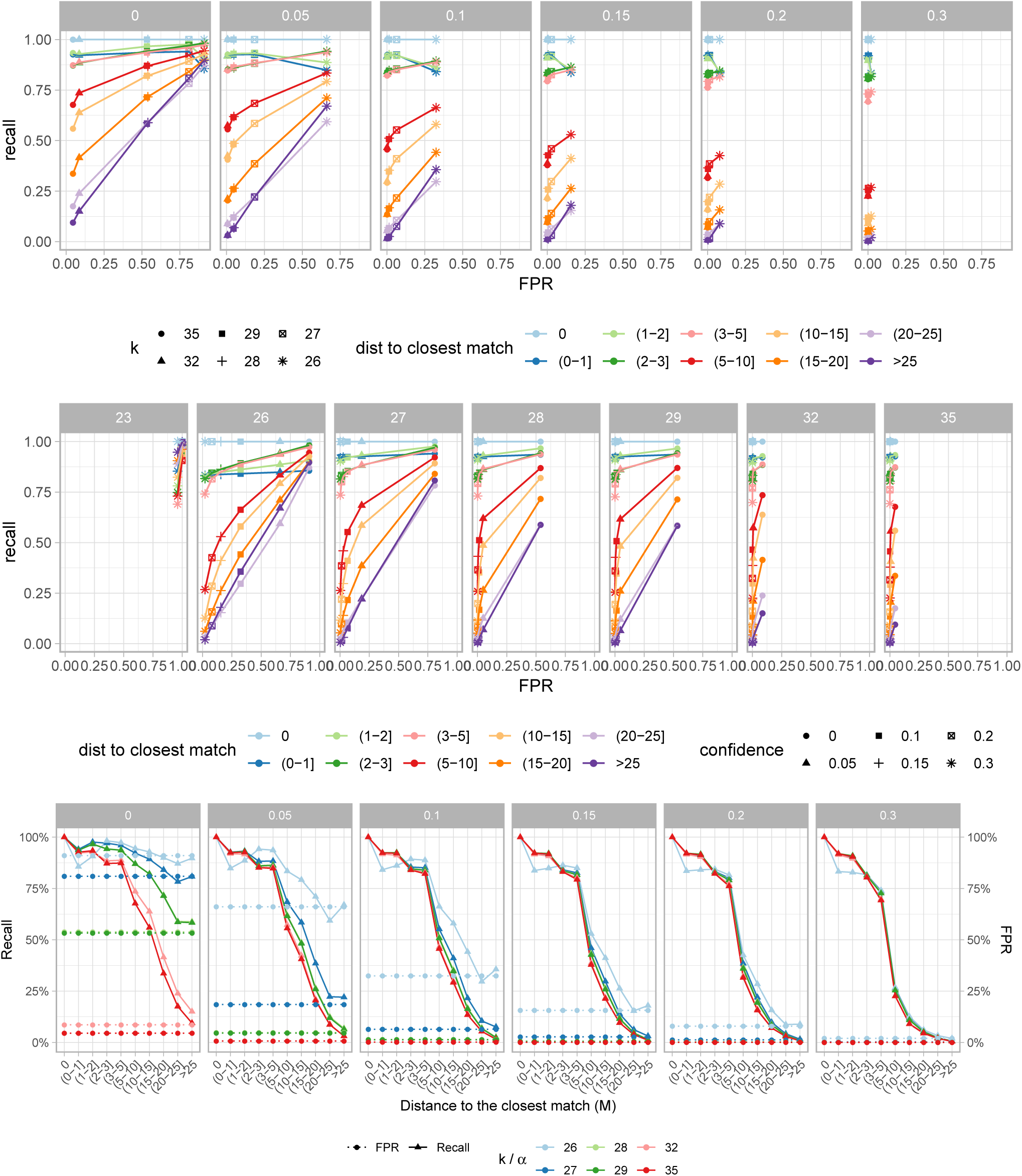
Roc.

**Figure S8:**
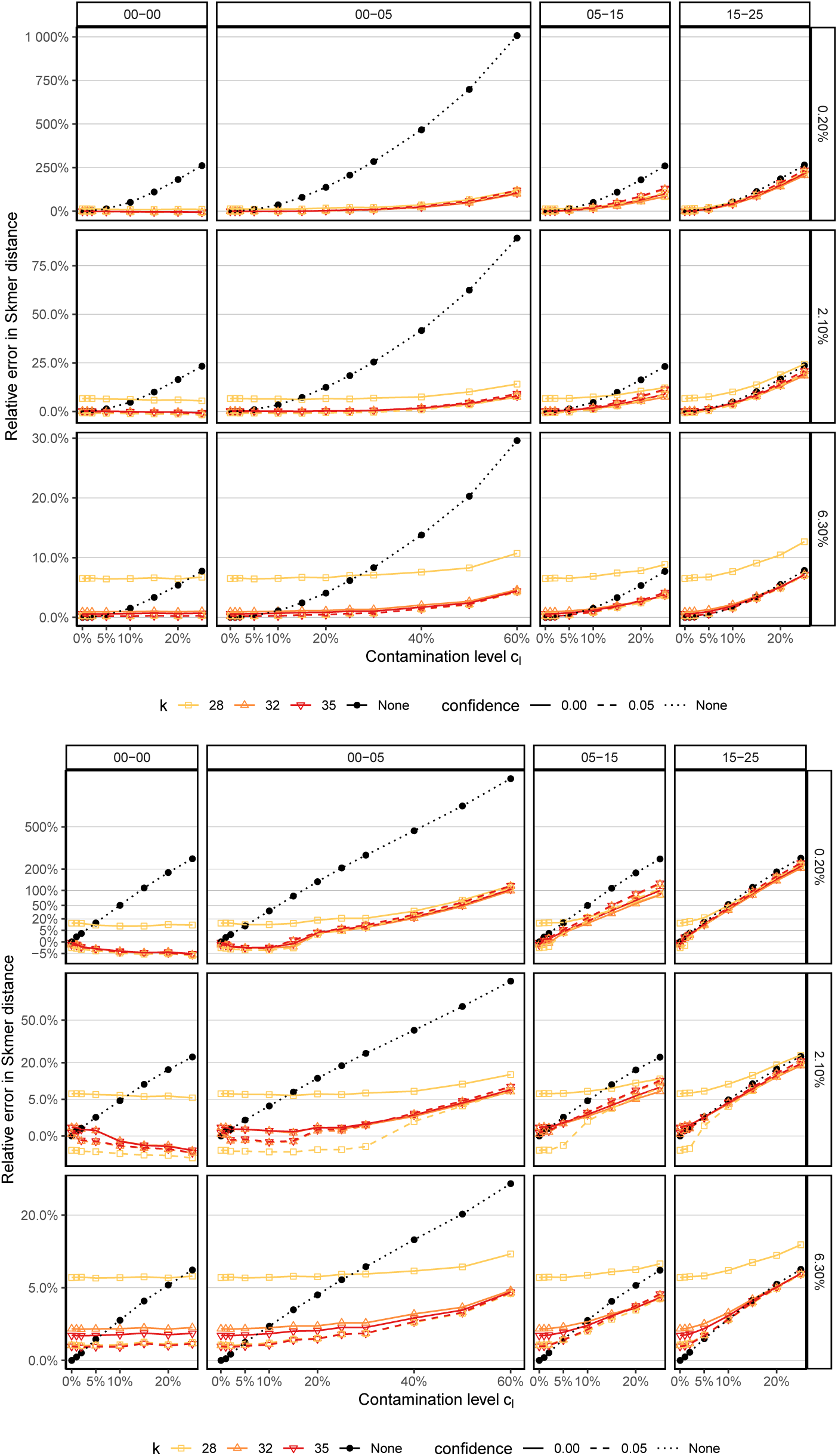
Relative error of Skmer distances without (None) and with (colored) Kraken-II filtering with different confidence levels *α* and *k*. Contaminants are added based on sequences that are at distance range *M* from the sequences in the reference library, for four ranges of *M* (boxes). The pairs of Drosophilas are chosen to be at true distance *D* = 0.2%, *D* = 2.1%, or *D* = 6.3%. top: normal scale; bottom: squre root scale.

**Figure S9:**
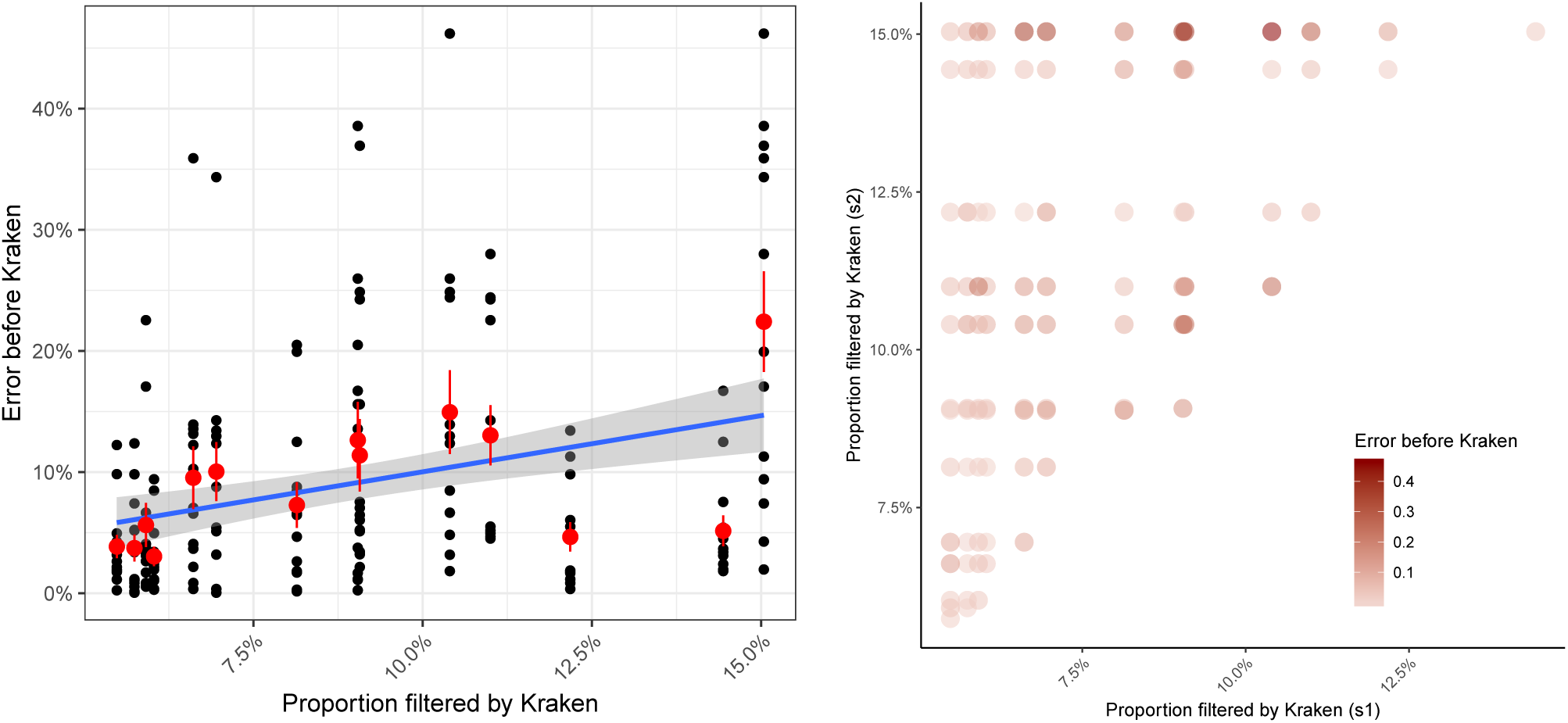
(a) Proportion of reads filtered by Kraken-II from one of the two species being compared versus the error before Kraken-II in the genomic distance. (b) For each pair of species, colors show the relative distance error before Kraken-II versus the proportion of reads filtered from each of the two genomes. Error is associated strongly, but imperfectly, with high levels of filtering.

**Figure S10:**
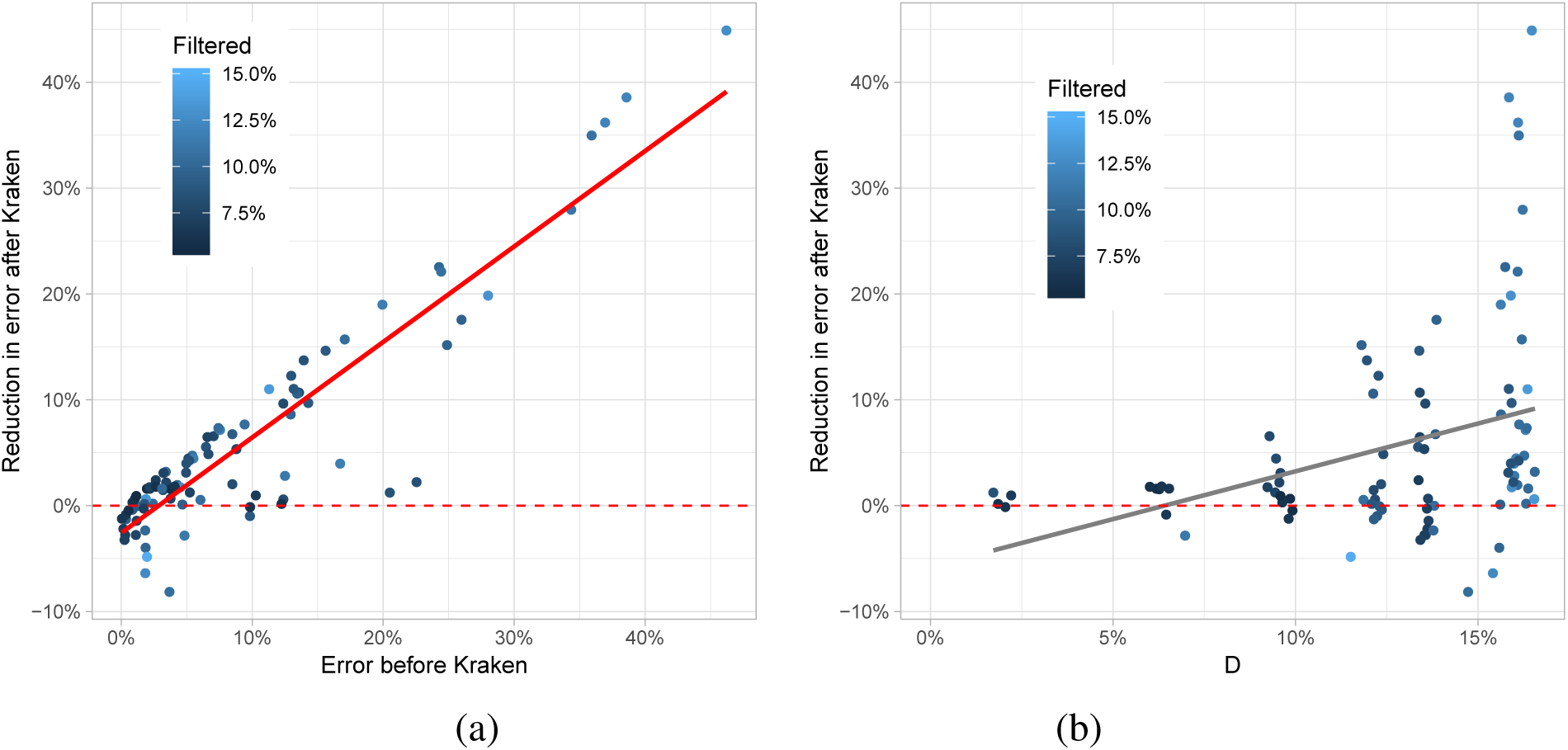
Change in relative distance error after filtering with Kraken-II for 100Mb Drosophila dataset versus (a) gold standard (assembly) genomic distance *D*, and (b) error before Kraken-II filtering.

**Figure S11:**
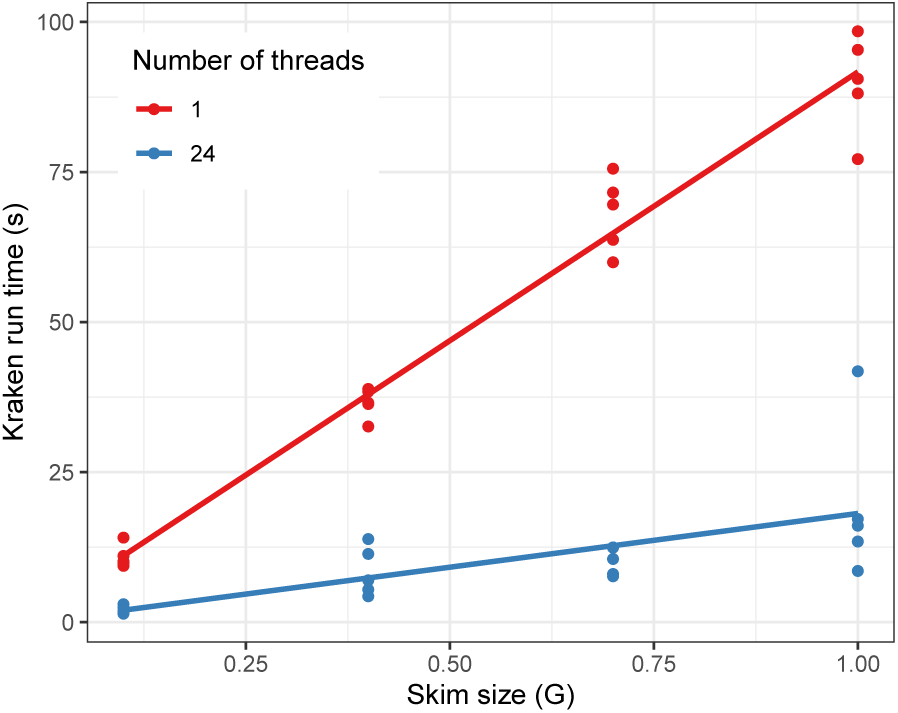
Kraken-II processing speed (s per query) with respect to different genome skim sizes. Sequences were simulated using ART (Huang et al., 2012) (same settings as before) to reach at least 1.4GB of synthetic reads and subsequently downsampled to generate 1GB, 0.7GB, 0.4GB and 0.1GB genome skim benchmark set. The dataset was queried using Kraken-II default settings and standard reference library. Kraken-II was run on a machine with Intel Xeon E5-2680v3 2.5 GHz CPU and 120GB of RAM running CentOS Linux release-6-10.

**Figure S12:**
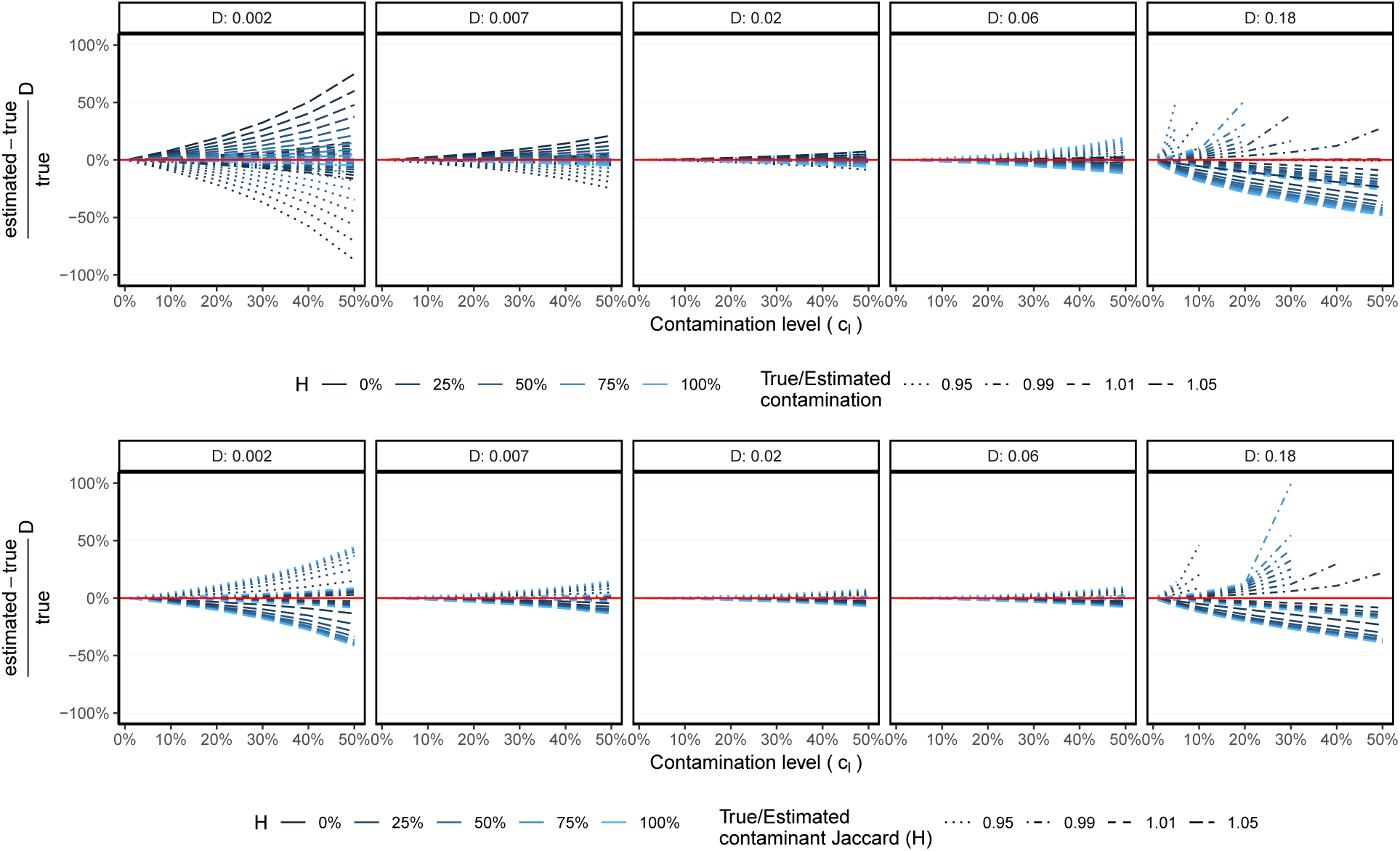
The sensitivity of filter-free correction using theoretical modelling. For six values of *D* (boxes) and varying *H*, relative error is shown for various contamination levels (setting *c*_*l*_ = *c*_1_ = *c*_2_) when the estimated distance is corrected using (6) when *c*_*l*_ is miscalculated by small margins (1% or 5% over or under-estimated). Missing values indicate cases where (6) gives undefined values. *k* = 31 in all cases.

## Appendix C Supplementary Tables

**Table S1:**
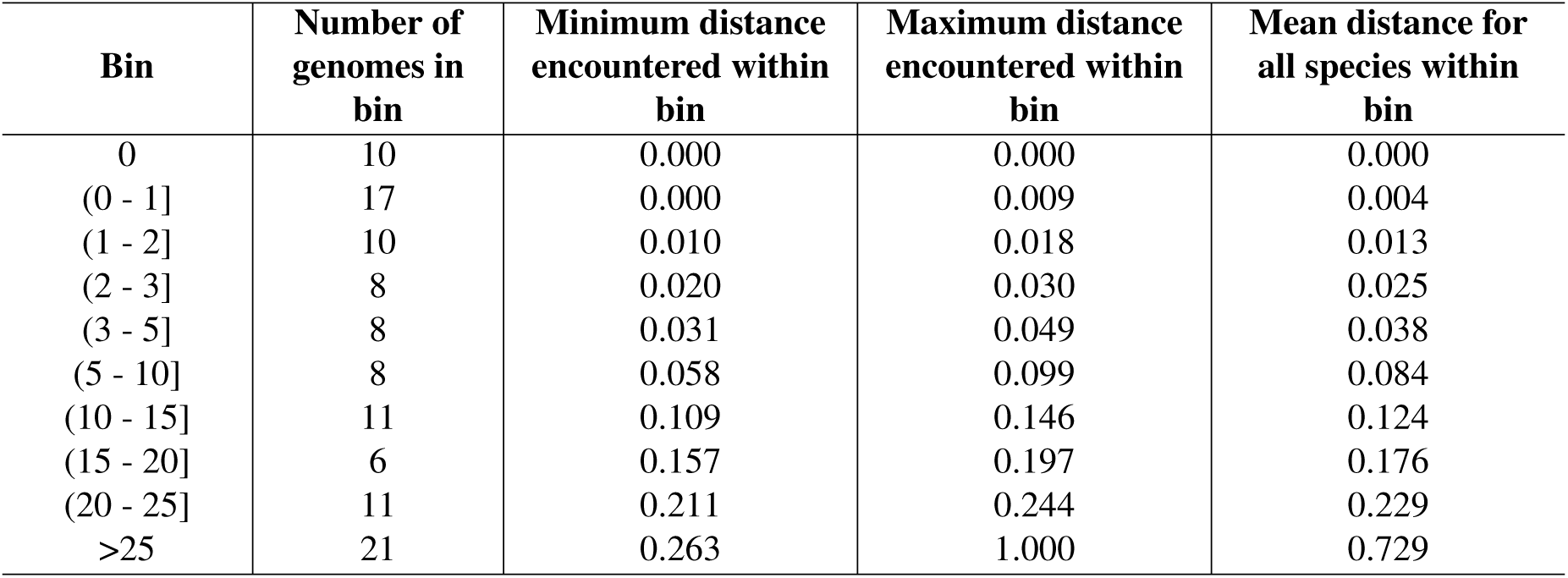
Bin assignment based on Mash distances.

**Table S2:** Plant species used as query sequences. Plants were selected to represent a wide range of genome sizes.

| Species name | Genome size (M) |
| --- | --- |
| Arabidopsis thaliana | 119.167 |
| Arabidopsis lyrata | 202.97 |
| Carya illinoensis | 649.75 |
| Carya cathayensis | 721.33 |
| Nicotiana sylvestris | 2221.99 |
| Zea mays | 2182.61 |
| Oryza sativa | 382.63 |
| Coffee arabica | 1094.45 |
| Prunus persica | 212.77 |
| Bathycoccus prasinos | 15.07 |

**Table S3:** Contamination level (*c*_*l*_) with the corresponding number of unique *k*-mers for base (*D. simulans w501*) and contaminant (bacteria) in mixture samples.

| Bin | Contamina-<br>tion level (%)<br>reads | $c_l$ (%): Unique<br>contaminant<br>$k$-mers in sample<br>(%) | Unique base $k$-mers<br>in sample (%) | Common $k$-mers<br>between base and<br>contaminant (%) |
| --- | --- | --- | --- | --- |
| 0-5 | 60 | 53.728625 | 46.271364 | 0.000011 |
| 0-5 | 50 | 46.000931 | 53.999065 | 0.000004 |
| 0-5 | 40 | 38.455505 | 61.544490 | 0.000005 |
| 0-5 | 30 | 30.730685 | 69.269315 | 0.000000 |
| 0-5 | 25 | 26.630857 | 73.369142 | 0.000001 |
| 0-5 | 20 | 22.286244 | 77.713750 | 0.000007 |
| 0-5 | 15 | 17.595401 | 82.404592 | 0.000007 |
| 0-5 | 10 | 12.408705 | 87.591295 | 0.000000 |
| 0-5 | 5 | 6.630852 | 93.369148 | 0.000000 |
| 0-5 | 2 | 2.769620 | 97.230380 | 0.000000 |
| 0-5 | 1 | 1.406449 | 98.593551 | 0.000000 |
| 0-5 | 0 | 0.000000 | 100.000000 | 0.000000 |
| 0-0 | 25 | 24.707830 | 75.292146 | 0.000024 |
| 0-0 | 20 | 20.899131 | 79.100838 | 0.000031 |
| 0-0 | 15 | 16.686621 | 83.313353 | 0.000026 |
| 0-0 | 10 | 11.968806 | 88.031170 | 0.000023 |
| 0-0 | 5 | 6.494518 | 93.505472 | 0.000010 |
| 0-0 | 2 | 2.742933 | 97.257067 | 0.000000 |
| 0-0 | 1 | 1.395767 | 98.604233 | 0.000000 |
| 0-0 | 0 | 0.000000 | 100.000000 | 0.000000 |
| 5-15 | 25 | 24.797418 | 75.202570 | 0.000012 |
| 5-15 | 20 | 20.977979 | 79.021978 | 0.000043 |
| 5-15 | 15 | 16.738171 | 83.261791 | 0.000038 |
| 5-15 | 10 | 12.010540 | 87.989451 | 0.000010 |
| 5-15 | 5 | 6.502650 | 93.497350 | 0.000000 |
| 5-15 | 2 | 2.745543 | 97.254457 | 0.000000 |
| 5-15 | 1 | 1.402008 | 98.597992 | 0.000000 |
| 5-15 | 0 | 0.000000 | 100.000000 | 0.000000 |
| 15-25 | 25 | 24.514304 | 75.485614 | 0.000082 |
| 15-25 | 20 | 20.760153 | 79.239792 | 0.000055 |
| 15-25 | 15 | 16.620725 | 83.379207 | 0.000068 |
| 15-25 | 10 | 11.916805 | 88.083179 | 0.000016 |
| 15-25 | 5 | 6.484925 | 93.515047 | 0.000028 |
| 15-25 | 2 | 2.738899 | 97.261078 | 0.000023 |
| 15-25 | 1 | 1.398807 | 98.601185 | 0.000009 |
| 15-25 | 0 | 0.000000 | 100.000000 | 0.000000 |

**Table S4:** Actual amount of non-unique *k*-mers (%) in contaminant reads used to generate simulated Drosophila skims with *H* overlap.

| Expected overlap (%) | 0.00 | 1.00 | 5.00 | 10.00 | 25.00 | 50.00 |
| --- | --- | --- | --- | --- | --- | --- |
| Actual $k$-mer overlap (%) | 11.12 | 11.54 | 13.31 | 15.83 | 23.82 | 40.78 |

**Table S5:**
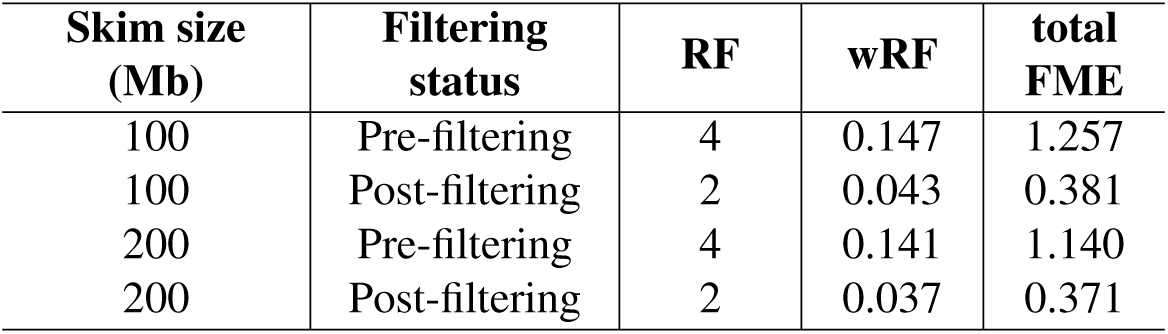
The effect of filtering on the quality of phylogenies inferred from genomic skims.

| **Skim size**<br>**(Mb)** | **Filtering**<br>**status** | **RF** | **wRF** | **total**<br>**FME** |
| --- | --- | --- | --- | --- |
| 100 | Pre-filtering | 4 | 0.147 | 1.257 |
| 100 | Post-filtering | 2 | 0.043 | 0.381 |
| 200 | Pre-filtering | 4 | 0.141 | 1.140 |
| 200 | Post-filtering | 2 | 0.037 | 0.371 |

**Table S6:**
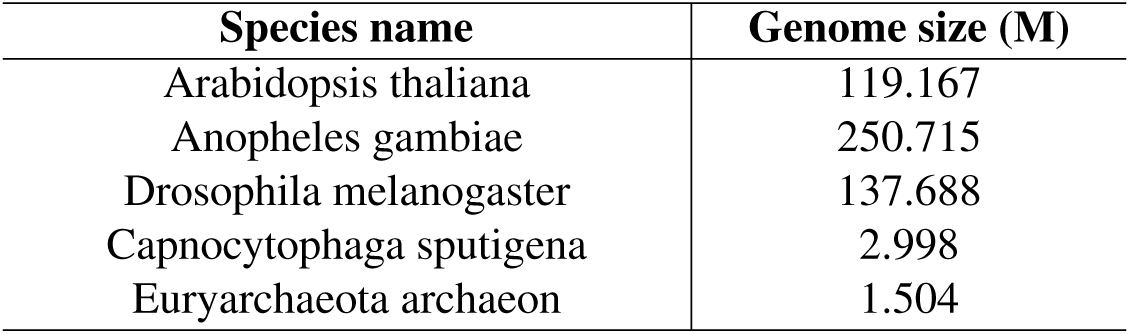
Species used as query sequences for assessing Kraken-II running time. Genomes were arbitrarily selected but represent a diverse set of both eukaryotic and prokaryotic species.

| **Species name** | **Genome size (M)** |
| --- | --- |
| Arabidopsis thaliana | 119.167 |
| Anopheles gambiae | 250.715 |
| Drosophila melanogaster | 137.688 |
| Capnocytophaga sputigena | 2.998 |
| Euryarchaeota archaeon | 1.504 |

## Appendix D Supplementary method details and commands

Here we provide the exact procedures and commands that we used to run external software throughout our experiments.

### Genome skims simulation

To simulate short reads with length *l* = 150 and coverage *c*, in single read mode with default error and quality profiles of Illumina HiSeq 2500, we ran

~~~
  art_illumina -ss HS25 -i FASTA_FILE -l 150 -f *c* -na -s 10 -o
FASTQ_FILE
~~~

### Downsampling reads

To subsample reads down a specified number of reads *n* we used

~~~
seqtk sample -s150 INPUT_FASTQ_FILE *n* > OUTPUT_FASTQ_FILE
~~~

### Reference library construction

To construct standard Kraken reference library we used default command

~~~
kraken2-build −−standard −−no-masking −−use-ftp −−db DATABASE_NAME
~~~

To build custom Kraken reference library we used a set of commands below:

- Download taxonomy

~~~
kraken2-build −−download-taxonomy −−no-masking −−use-ftp −−db
DATABASE_NAME
~~~
- Download viral database

~~~
kraken2-build −−download-library viral −−no-masking −−use-ftp
−−db DATABASE_NAME
~~~
- Rename file extensions to .fa

~~~
find. -name “*.fna” -exec sh -c ‘mv “$1” “${1%.fna}.fa”’ _ {} \;
~~~
- Add custom genomes to the reference library

~~~
find genomes/ -name ‘*.fa’ -print0 | xargs -0 -I -n1 kraken2-build
−−no-masking −−add-to-library {} −−db DATABASE_NAME
~~~
- Build database with specified *k*-mer length *k*, minimizer length *l* and number of wind-carding positions *s* we used

~~~
kraken2-build −−build −−no-masking −−kmer-len *k* −−minimizer-len *l*
−−minimizer-spaces *s* −−use-ftp −−db DATABASE_NAME
~~~

### Reference library querying

To query reference library at variable confidence level *α* we used

~~~
  kraken2 −−use-names −−threads 24 −−report REPORT_FILE_NAME −−db
DATABASE_NAME −−confidence *α* −−classified-out
CLASSIFIED_FASTQ_FILE −−unclassified-out UNCLASSIFIED_FASTQ_FILE
QUERY_FASTQ_FILE > KRAKEN_OUTPUT_FILE
~~~

### Computing k-mer frequencies

To estimate *k*-mer frequencies we used Jellyfish.

- Computing *k*-mer profile

~~~
jellyfish count -m 31 -s 100M -t 18 -C INPUT_FASTQ_FILE
-o COUNT_FASTQ_FILE
~~~
- Extracting *k*-mer statistics

~~~
jellyfish stats COUNT_FASTQ_FILE
~~~

### Computing genomic distances

To estimate genomic distances we used Mash and Skmer.

- To compute genomic distance using Mash we used

~~~
mash dist FASTQ_FILE_ONE FASTQ_FILE_TWO
~~~
- To compute genomic distance using Skmer we used

~~~
skmer reference FASTQ_DIRECTORY -p 24 -o REF_DISTANCE_MATRIX
~~~

### Preprocessing real data sequencing files

We used BBTools to preprocess real data sequencing read files.

- To decontaminate .fastq files we used

~~~
bbduk.sh in1=FASTQ_READ1 in2=FASTQ_READ2 out1=FASTQ_READ1 out2=FASTQ_READ2
ref=adapters,phix ktrim=r k=23 mink=11 hdist=1 tpe tbo
~~~
- To deduplicate reads we used

~~~
dedupe.sh in1=FASTQ_READ1 in2=FASTQ_READ2 out=DEDUP_OUTPUT_FASTQ_FILE
~~~
- To reformat deduplicated output files we used

~~~
reformat.sh in=DEDUP_OUTPUT_FASTQ_FILE out1=FASTQ_READ1
out2=FASTQ_READ2
~~~
- To merge paired-end reads .fastq we used

~~~
bbmerge.sh in1=FASTQ_READ1 in2=FASTQ_READ2 out1=OUTPUT_FASTQ_FILE
~~~

## Appendix E Raw data

**Table S7:**
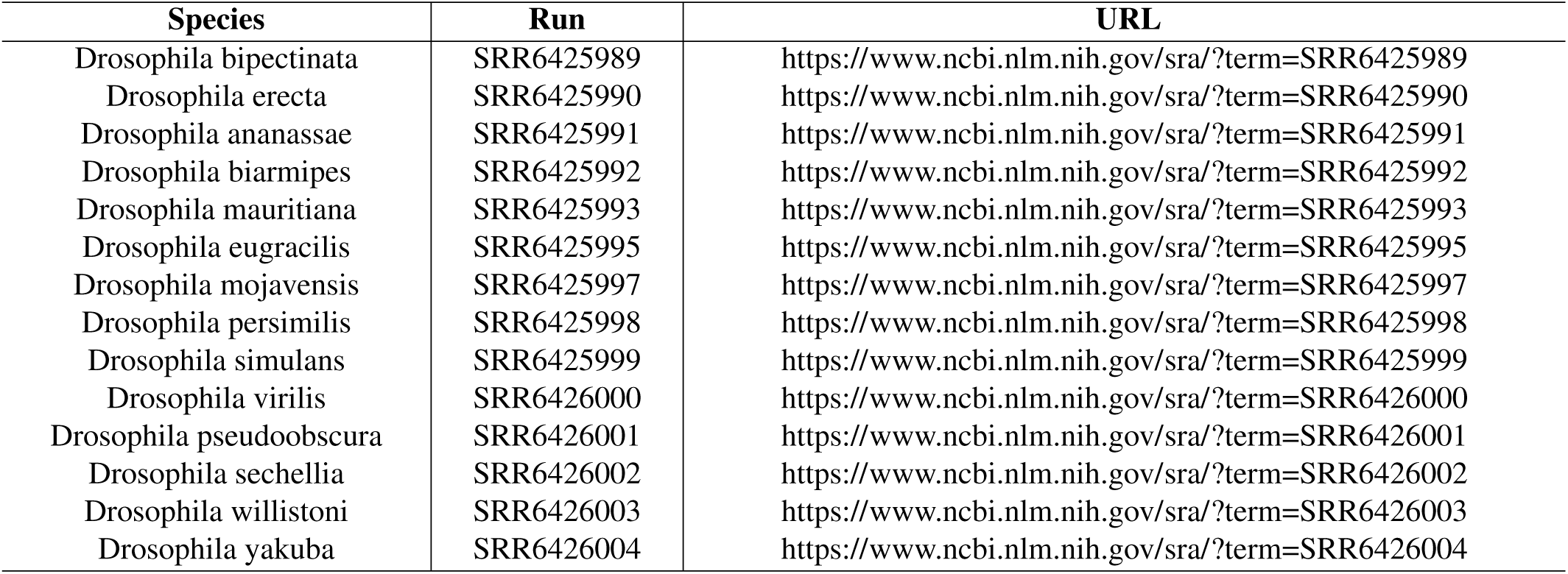
List of SRRs and URLs for Drosophila species used in real data experiment.

**Table S8:** List of accession numbers and URLs for Drosophila species used in contamination simulation experiment.

| Species | Assembly accession | URL |
| --- | --- | --- |
| *Drosophila simulans* w501 | GCF_000754195.2 | https://www.ncbi.nlm.nih.gov/assembly/GCF_000754195.2 |
| *Drosophila simulans* WXD1 | GCA_004382185.1 | https://www.ncbi.nlm.nih.gov/assembly/GCA_004382185.1 |
| *Drosophila sechellia* | GCF_000005215.3 | https://www.ncbi.nlm.nih.gov/assembly/GCF_000005215.3 |
| *Drosophila yakuba* | GCF_000005975.2 | https://www.ncbi.nlm.nih.gov/assembly/GCF_000005975.2 |

**Table S9:**
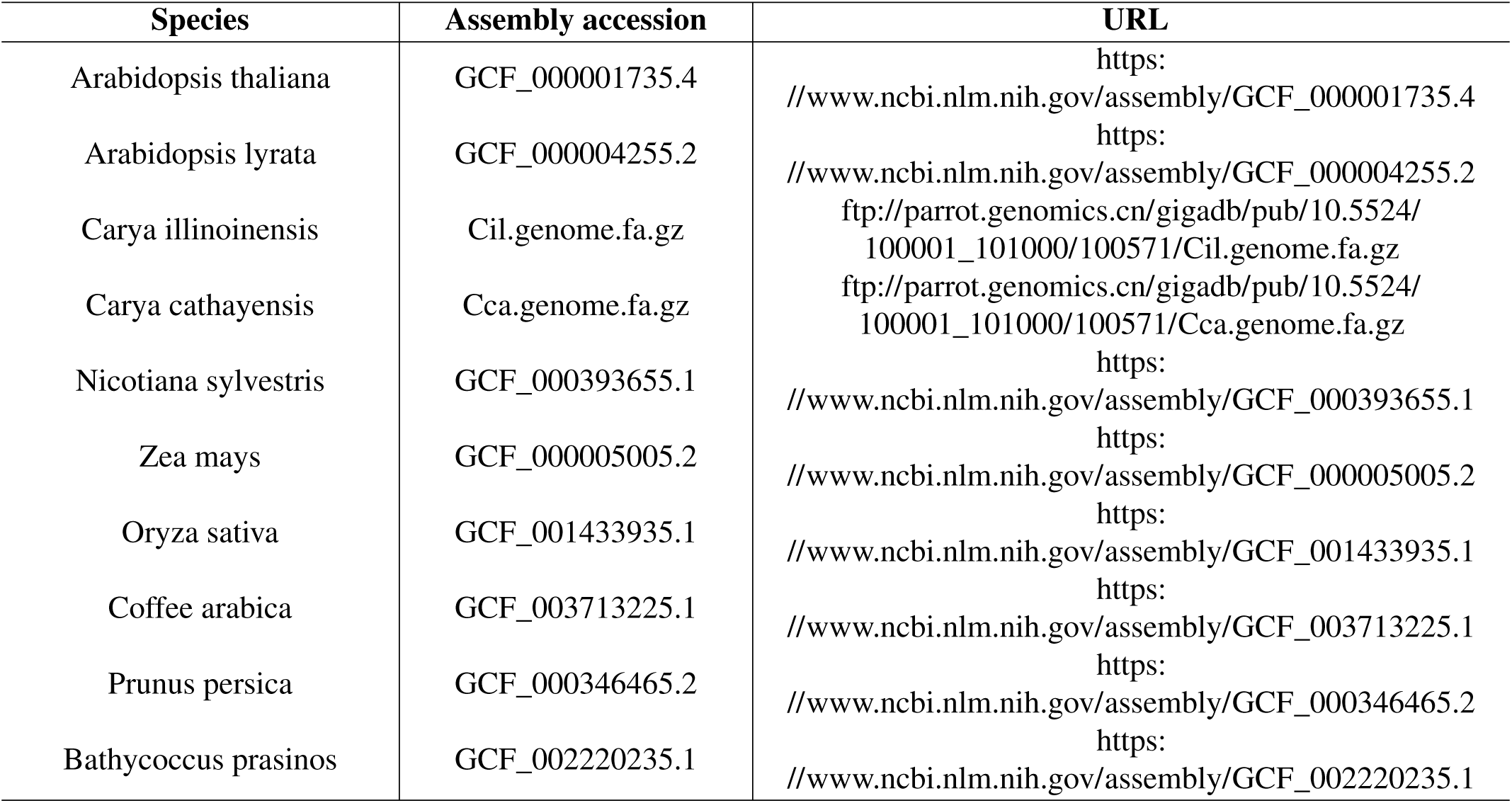
List of accession numbers and URLs for plant species added to query set.

| Species | Assembly accession | URL |
| --- | --- | --- |
| Arabidopsis thaliana | GCF_000001735.4 | https:<br>//www.ncbi.nlm.nih.gov/assembly/GCF_000001735.4 |
| Arabidopsis lyrata | GCF_000004255.2 | https:<br>//www.ncbi.nlm.nih.gov/assembly/GCF_000004255.2 |
| Carya illinoensis | Cil.genome.fa.gz | ftp://parrot.genomics.cn/gigadb/pub/10.5524/<br>100001_101000/100571/Cil.genome.fa.gz |
| Carya cathayensis | Cca.genome.fa.gz | ftp://parrot.genomics.cn/gigadb/pub/10.5524/<br>100001_101000/100571/Cca.genome.fa.gz |
| Nicotiana sylvestris | GCF_000393655.1 | https:<br>//www.ncbi.nlm.nih.gov/assembly/GCF_000393655.1 |
| Zea mays | GCF_000005005.2 | https:<br>//www.ncbi.nlm.nih.gov/assembly/GCF_000005005.2 |
| Oryza sativa | GCF_001433935.1 | https:<br>//www.ncbi.nlm.nih.gov/assembly/GCF_001433935.1 |
| Coffee arabica | GCF_003713225.1 | https:<br>//www.ncbi.nlm.nih.gov/assembly/GCF_003713225.1 |
| Prunus persica | GCF_000346465.2 | https:<br>//www.ncbi.nlm.nih.gov/assembly/GCF_000346465.2 |
| Bathycoccus prasinos | GCF_002220235.1 | https:<br>//www.ncbi.nlm.nih.gov/assembly/GCF_002220235.1 |

